# Dissection of the Spindle Assembly Checkpoint by Proximity Proteomics

**DOI:** 10.1101/2020.06.04.133710

**Authors:** Yenni A. Garcia, Erick F. Velasquez, Lucy W. Gao, Keith Cheung, Kevin M. Clutario, Taylor Williams-Hamilton, Ankur A. Gholkar, Julian P. Whitelegge, Jorge Z. Torres

## Abstract

The spindle assembly checkpoint (SAC) is critical for sensing defective microtubule-kinetochore attachments and tension across the kinetochore and functions to arrest cells in prometaphase to allow time to repair any errors prior to proceeding into anaphase. The SAC has a central role in ensuring the fidelity of chromosome segregation and its dysregulation has been linked to the development of human diseases like cancer. The establishment and maintenance of the SAC relies on multiple protein complexes that are intricately regulated in a spatial and temporal manner through posttranslational modifications like phosphorylation. Over the past few decades the SAC has been highly investigated and much has been learned about its protein constituents and the pathways and factors that regulate its activity. However, the spatio-temporal proximity associations of the core SAC components have not been explored in a systematic manner. Here, we have taken a BioID2 proximity-labeling proteomic approach to define the proximity protein environment for each of the five core SAC proteins BUB1, BUB3, BUBR1, MAD1L1, and MAD2L1 under conditions where the SAC is active in prometaphase. These five protein association maps were integrated to generate the SAC proximity protein network that contains multiple layers of information related to core SAC protein complexes, protein-protein interactions, and proximity associations. Our analysis validated many of the known SAC complexes and protein-protein interactions. Additionally, it uncovered new protein associations that lend insight into the functioning of the SAC and highlighted future areas that should be investigated to generate a comprehensive understanding of the SAC.

## INTRODUCTION

Human cell division is a highly coordinated set of events that ensures the proper transmission of genetic material from one mother cell to two newly formed daughter cells. Chromosome missegregation during cell division can lead to aneuploidy, an aberrant chromosomal number, which is a hallmark of many types of cancers and has been proposed to promote tumorigenesis (1). However, there is currently no consensus as to the pathways and factors that are deregulated to induce aneuploidy, why it is prevalent in cancer and how it contributes to tumorigenesis. Pivotal to cell division is the metaphase to anaphase transition, which is a particularly regulated process involving a multitude of protein-protein interactions that relies heavily on posttranslational modifications like phosphorylation and ubiquitination that function as switches to activate or inactivate protein function (2,3). For example, the multi-component spindle assembly checkpoint (SAC) is activated when unattached kinetochores or nonproductive (monotelic, syntelic, and merotelic) attachments are sensed and functions to arrest cells in metaphase to give time to correct these deficiencies and generate proper microtubule-kinetochore attachments (2) (Fig. 1*A*). This ensures proper sister chromatid separation and minimizes segregation errors that lead to chromosomal instability, aneuploidy, and tumorigenesis (1). Core components of the SAC include BUB1, BUB3, BUBR1, MAD1L1, and MAD2L1(4). Critical to the SAC is the mitotic checkpoint complex (MCC, composed of MAD2L1, BUBR1, BUB3, and CDC20) that maintains the anaphase promoting complex/cyclosome (APC/C) ubiquitin ligase substrate adaptor protein CDC20 sequestered and thereby inactivates the APC/C (5,6). Upon proper microtubule-kinetochore attachment the SAC is satisfied and the inhibitory effect of the MCC on the APC/C is relieved (2) (Fig. 1*A*). Active APC/C then ubiquitinates and targets Securin for degradation (2), which activates Separase, the protease that cleaves RAD21, a component of the cohesin complex that holds sister chromatids together (7). This releases sister chromatid cohesion and chromatids are pulled to opposing poles of the cell by spindle microtubules, marking the entry into anaphase.

**Fig. 1.**
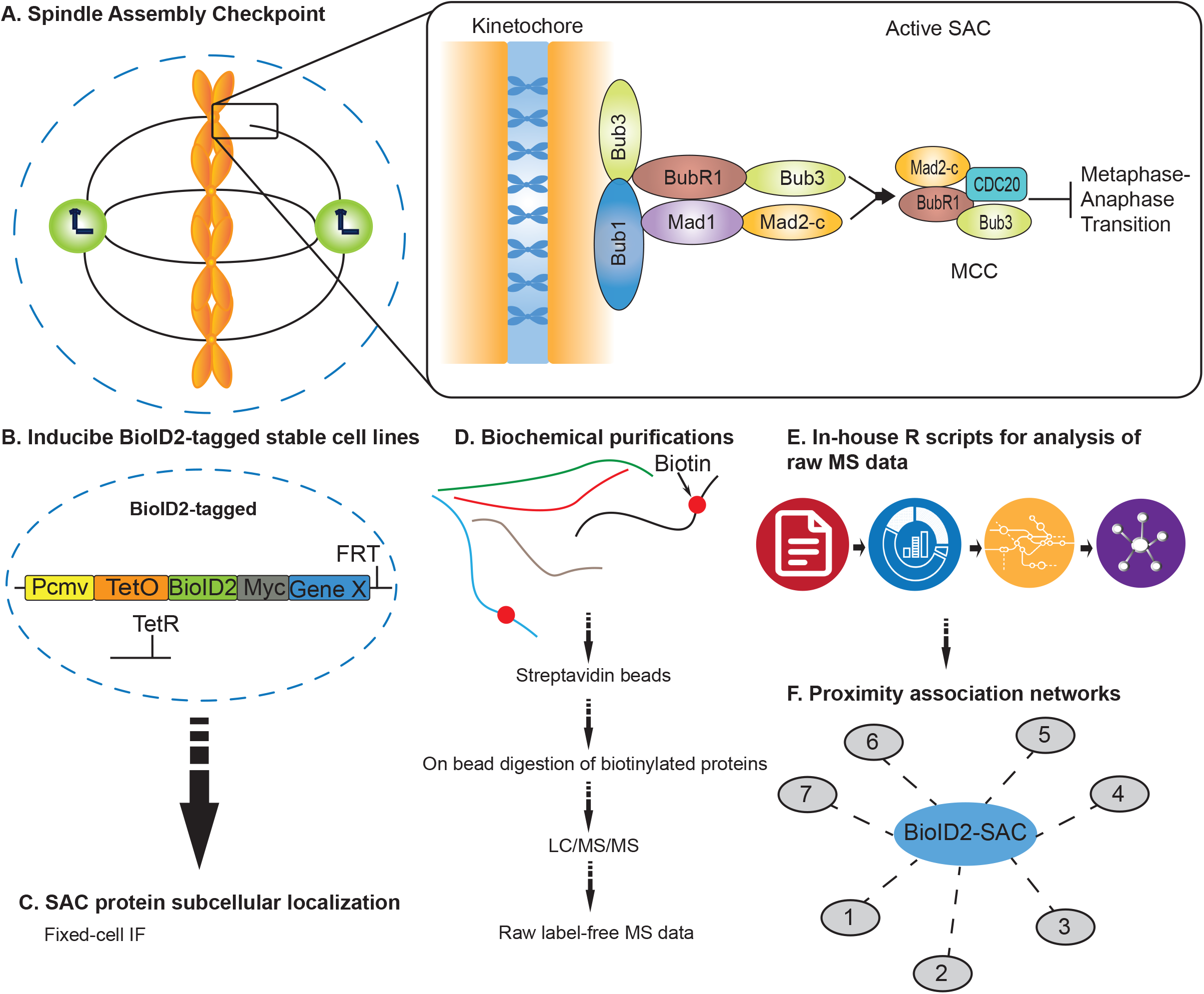
Overview of the approach to generate core SAC protein BioID2 proximity association networks. *A*, Schematic of the core spindle assembly checkpoint (SAC) components BUB1, BUB3, BUBR1, MAD1L1, and MAD2L1 that localize to the kinetochore region during early mitosis. MCC denotes mitotic checkpoint complex. *B*, Generation of inducible BioID2-tagged stable cell lines for each core SAC protein. *C*, Fixed-cell immunofluorescence microscopy to analyze BioID2-tagged SAC protein subcellular localization in time and space. *D*, Biochemical purifications; affinity purification of biotinylated proteins and identification of proteins by LC/MS/MS. *E*, Computational analysis of raw mass spectrometry data using in-house R scripts. *F*, Generation of high-confidence SAC protein proximity association networks.

Because understanding the SAC is critical to understanding tumorigenesis and the response of tumor cells to antimitotic drugs that activate the SAC and trigger apoptotic cell death, it has become an intensive area of research (8,9). Although decades of research have shed light on the SAC, we are far from elucidating the full complement of regulatory factors involved in this complex pathway and from understanding how misregulation of this pathway can lead to tumorigenesis and resistance to chemotherapeutic drugs like antimitotics (10). Furthermore, models of the spatial temporal associations of the core SAC proteins with themselves and with structural and signaling components that mediate the establishment and silencing of the SAC are still being defined (11–13). Recently, proximity-labeling approaches like BioID and APEX have been used effectively to determine the spatial and temporal associations among proteins and for defining the architecture of centrosome, centrosome-cilia interface, and other organelles within the cell (14–19). However, proximity labeling has not been applied to the SAC in a systematic fashion, which could help to interrogate current models of core SAC protein associations and regulation.

Here, we have engineered vectors for establishing inducible BioID2-tagged protein stable cell lines. This system was used to establish stable cell lines with inducible BioID2-tagged core SAC protein (BUB1, BUB3, BUBR1, MAD1L1, and MAD2L1) expression. These cell lines were utilized in BioID2-proximity biotin labeling studies, which were coupled to biotin biochemical purifications and mass spectrometry analyses to map the spatial and temporal associations among the core SAC proteins and other proteins in close proximity. These analyses yielded a wealth of information with regards to the protein environment of the core SAC proteins under conditions where the SAC is active. In addition to validating well-established SAC protein complexes and protein-protein interactions, we defined new protein associations that advance our understanding of the SAC and we highlight areas of research that warrant further investigation to further comprehend the SAC and its regulation.

## EXPERIMENTAL PROCEDURES

### Cell Culture and Cell Cycle Synchronization

All media and chemicals were purchased from ThermoFisher Scientific (Waltham, MA) unless otherwise noted. HeLa Flp-In T-REx BioID2-tagged stable cell lines and RPE cells were grown in F12:DMEM 50:50 medium with 10% FBS, 2 mM L-glutamine and antibiotics, in 5% CO_2_ at 37° C. Cells were induced to express the indicated BioID2-tagged proteins by the addition of 0.2 μg/ml doxycycline (Sigma-Aldrich, St. Louis, MO) for 16 hours. For synchronization of cells in mitosis, cells were treated with 100 nM Taxol (Sigma-Aldrich) for 16 hours. For a list of all reagents used see supplemental Table S1.

### Cell siRNA and Chemical Treatments

HeLa cell siRNA treatments were performed as described previously (20), with control siRNA (siControl, D-001810-10) or BUB1-targeting siRNA (siBUB1, L-004102-00) from Dharmacon (Lafayette, CO) for 48 hours. For chemical treatments, RPE or HeLa cells were treated with control DMSO vehicle or the BUB1 inhibitor BAY 1816032 (HY-103020) (21) from MedChemExpress (Monmouth Junction, NJ) at 10 μM for five hours.

### Generation of Inducible BioID2-tagging Vectors and Stable Cell Lines

For generating pGBioID2-27 or pGBioID2-47 vectors, the EGFP-S-tag was removed from pGLAP1 (22) by digestion with BstBI and AflII. BioID2-Myc-27 (27 amino acid linker) or BioID2-Myc-47 (47 amino acid linker) were PCR amplified, digested with NheI and XhoI and cloned into BstBI and AflII digested pGLAP1 to generate pGBioID2-27 or pGBioID2-47 (supplemental Fig. S1*A*). For full-length human SAC core gene *hBUB1, hBUB3, hBUBR1, hMAD1L1,* and *hMAD2L1* expression, cDNA corresponding to the full-length open reading frame of each gene was cloned into pDONR221 as described previously (22,23) (supplemental Fig. S1*B*). SAC core genes were then transferred from pDONR221 to pGBioID2-47 using the Gateway cloning system (Invitrogen, Carlsbad, CA) as described previously (22,23) (supplemental Fig. S1*B*). The pGBioID2-47-SAC protein vectors were then used to generate doxycycline inducible HeLa Flp-In T-REx BioID2 stable cell lines that expressed the fusion proteins from a specific single locus within the genome as described previously (22,23) (supplemental Fig. S1*C* and 1*D*). All primers were purchased from ThermoFisher Scientific. For a list of primers used see supplemental Table S2. For a list of vectors generated in this study see supplemental Table S3. The pGBioID2-27 and pGBioID2-47 vectors have been deposited at Addgene (AddgeneIDs: 140276 and 140277 respectively) and are available to the scientific community.

### Biotin Affinity Purifications

All media, chemicals, and beads were purchased from ThermoFisher Scientific unless otherwise noted. Biotin affinity purifications were conducted using previously described protocols with modifications (18,19). Briefly, 10% FBS was treated with 1 ml of MyOne Streptavidin C1 Dynabeads overnight and passed through a 0.22 μm filter. The BioID2-BUB1, BUB3, BUBR1, MAD1L1, and MAD2L1, and BioID2 alone inducible stable cell lines were plated on six 150 mm tissue culture dishes, 24 hours post-plating, the cells were washed three times with PBS and once with DMEM without FBS, and shifted to the streptavidin Dynabead-treated 10% FBS DMEM. The cells were induced with 0.2 μg/ml Dox, and treated with 100 nM Taxol and 50 mM Biotin for 16 hours. Mitotic cells were collected and centrifuged at 1,500 rpm for 5 minutes and washed twice with PBS. The pellet was lysed with 3 ml of lysis buffer (50 mM Tris-HCl pH 7.5, 150 mM NaCl, 1 mM EDTA, 1 mM EGTA, 1% Triton-X-100, 0.1% SDS, Halt Protease and Phosphatase Inhibitor Cocktail) and incubated with gentle rotation for 1 hour at 4° C, then centrifuged at 15,000 rpm for 15 minutes and transferred to a new 15 ml conical tube. The lysate was transferred to a TLA-100.3 tube (Beckman Coulter, Indianapolis, IN) and centrifuged at 45,000 rpm for 1 hour at 4° C. The lysate was then transferred to a new 15 ml conical tube and incubated with 300 μl of equilibrated streptavidin Dynabeads overnight with gentle rotation at 4° C. The beads were separated with a magnetic stand and washed twice with 2% SDS, followed by a wash with WB1 (0.1% sodium deoxycholate, 1% Triton X-100, 500 mM NaCl, 1 mM EDTA, 50 mM HEPES), a wash with WB2 (250 mM LiCl, 0.5% deoxycholate, 1 mM EDTA, 10 mM Tris-HCl pH 8.0), and a final wash with 50 mM Tris-HCl pH 7.5. The beads were then resuspended in 50 mM triethylammonium bicarbonate (TEAB), 12 mM sodium lauroyl sarcosine, 0.5% sodium deoxycholate. 10% of the beads were boiled with sample buffer and used for immunoblot analysis.

### In Solution Tryptic Digestion

Streptavidin Dynabeads in 50 mM triethylammonium bicarbonate (TEAB), 12 mM sodium lauroyl sarcosine, 0.5% sodium deoxycholate were heated to 95° C for 10 minutes and then sonicated for 10 minutes to denature proteins. Protein disulfide bonds were reduced by treatment with 5 mM tris(2-carboxyethyl) phosphine (final concentration) for 30 minutes at 37° C. Protein alkylation was performed with 10 mM chloroacetamide (final concentration) and incubation in the dark for 30 minutes at room temperature. The protein solutions were diluted five-fold with 50 mM TEAB. Trypsin was prepared in 50 mM TEAB and added 1:100 (mass:mass) ratio to target proteins followed by a 4-hour incubation at 37° C. Trypsin was again prepared in 50 mM TEAB and added 1:100 (mass:mass) ratio to target proteins followed by overnight incubation at 37° C. A 1:1 (volume:volume) ratio of ethyl acetate plus 1% trifluoroacetic acid (TFA) was added to the samples and samples were vortexed for five minutes. Samples were centrifuged at 16,000 x *g* for five minutes at room temperature and the supernatant was discarded. Samples were then lyophilized by SpeedVac (ThermoFisher Scientific) and desalted on C18 StageTips (ThermoFisher Scientific) as described previously (24).

### Nano-liquid Chromatography with Tandem Mass Spectrometry (LC-MS/MS) Analysis

Nano-LC-MS/MS with collision-induced dissociation was performed on a Q Exactive Plus Orbitrap (ThermoFisher Scientific) integrated with an Eksigent 2D nano-LC instrument, as described previously (25).

### Experimental Design and Statistical Rationale

To enhance confidence in identifying core SAC protein proximity associations, we performed control and experimental purifications in biological replicates (n=3 biological purifications for all core SAC proteins (except for BUB3 where n=2 biological purifications) and n=2 technical replicates each). This approach allowed for downstream comparison of control and experimental purifications, where proteins identified in the control BirA only (empty vector) were deemed potential non-specific associations. Database searches of the acquired spectra were analyzed with Mascot (v2.4; Matrix Science, Boston, MA) as described previously (25). The UniProt human database (October 10, 2018) was used with the following search parameters: trypsin digestion allowing up to 2 missed cleavages, carbamidomethyl on cysteine as a fixed modification, oxidation of methionine as a variable modification, 10-ppm peptide mass tolerance, and 0.5-Da fragment mass tolerance. With these parameters, an overall 5% peptide false discovery rate, which accounts for total false positives and false negatives, was obtained using the reverse UniProt human database as the decoy database. Peptides that surpassed an expectation cut-off score of 20 were accepted. All raw mass spectrometry files can be accessed at the UCSD Center for Computational Mass Spectrometry MassIVE datasets ftp://MSV000084975@massive.ucsd.edu (login: Torres, password: mitosis1). Peptides meeting the above criteria with information about their corresponding identified protein were further analyzed using in-house R scripts. All R scripts used in this study are freely available at GitHub https://github.com/torreslabucla/SpindleAssemblyBioID/. To increase precision and reduce error, a pseudo qualitative/quantitative approach was taken. Proteins identified in both the control and test purifications were assayed for significance, whereas proteins identified in test purifications but not present in control purifications were further considered. To handle proteins shared between test and control purifications, but only identified in less frequency, we measured the relative fold change or mean difference in a quantitative manner. To compare quantification between purifications, we used the Exponentially Modified Protein Abundance Index (emPAI) (26). emPAI offers approximate relative quantitation of the proteins in a mixture based on protein coverage by the peptide matches in a database search result and can be calculated using the following equation (26).

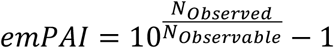

Where N_Observed_ is the number of experimentally observed peptides and N_Observable_ is the calculated number of observable peptides for each protein (26). Using emPAI as a relative quantification score, we calculated the mean difference (the mean emPAI for a certain protein across test replicates minus the mean emPAI for the same protein across control purifications). Using resampling in a paired manner, we then estimated the distribution of the mean difference for shared proteins between test and control purifications (27). Resampling involved recreating or estimating the normal distribution around a test statistic, in this case the mean difference, by calculating that statistic many times under rearrangement of labels. We performed ten thousand simulations per test statistic, resulting in normal distributions of mean difference between values of proteins identified in the experimental and the control. Using this distribution, we related each individual mean difference to the mean difference observed in the overall population in order to get a relative idea of what might be significantly higher in value compared to the control, when taking what is observed in the entire population. Values that lied outside of the 95% confidence interval of the mean difference and showed a higher value in the experimental compared to the control were then considered for further analysis (see supplemental Table S4).

### Protein Proximity Network Visualization and Integration of Systems Biology Databases

Visual renderings relating protein-protein interactions/associations were carried out using custom scripts in R. To incorporate protein-complex information, we integrated the Comprehensive Resource of Mammalian Protein Complexes (CORUM v. 3.0) (28). Protein-protein interaction information was derived and integrated from the Biological General Repository for Interaction Datasets (BioGRID v. 3.5) (29). To create relational networks that associated proteins based on cellular mechanisms, Gene Ontology (GO) terms were incorporated into the search (Gene Ontology release June 2019) (30). For a list of GO terms used, see supplemental Table S5. Pathway information was derived from Reactome, an open source and peer-reviewed pathway database (31). All databases were individually curated into an in-house systems biology relational database using custom R scripts. Final visuals relating protein associations were constructed using RCytoscapeJS, a developmental tool used to develop Cytoscape renderings in an R and JavaScript environment (32,33).

### Immunofluorescence Microscopy

Immunofluorescence microscopy was performed as described previously (34) with modifications described in (25). Briefly, HeLa inducible BioID2-tagged BUB1, BUB3, BUBR1, MAD1L1, and MAD2L1 stable cell lines were treated with 0.2 μg/ml doxycycline for 16 hours, fixed with 4% paraformaldehyde, permeabilized with 0.2% Triton X-100/PBS, and co-stained with 0.5 μg/ml Hoechst 33342 and the indicated antibodies. Imaging of mitotic cells was carried out with a Leica DMI6000 microscope (Leica DFC360 FX Camera, 63x/1.40-0.60 NA oil objective, Leica AF6000 software, Buffalo Grove, IL) at room temperature. Images were subjected to Leica Application Suite 3D Deconvolution software and exported as TIFF files.

### Antibodies

Immunofluorescence microscopy and immunoblotting were performed using the following antibodies: BioID2 (BioFront Technologies, Tallahassee, FL), GAPDH (Preoteintech, Rosemont, IL), α-tubulin (Serotec, Raleigh, NC), anti-centromere antibody (ACA, Cortex Biochem, Concord, MA), SGO2 (Bethyl, Montgomery, TX), PLK1 (Abcam, Cambridge, MA). Affinipure secondary antibodies labeled with FITC, Cy3, and Cy5 were purchased from Jackson Immuno Research (West Grove, PA). Immunoblot analyses were carried out using secondary antibodies conjugated to IRDye 680 and IRDye 800 from LI-COR Biosciences (Lincoln, NE) and blots were scanned using a LI-COR Odyssey infrared imager.

## RESULTS

### Generation of Inducible BioID2-tagged SAC Protein Stable Cell Lines

The spindle assembly checkpoint is essential for ensuring the fidelity of chromosome segregation during cell division (35) (Fig. 1*A*). To better understand how the SAC functions and is regulated in a spatial and temporal manner, we sought to map the protein associations of the core SAC proteins BUB1, BUB3, BUBR1 (BUB1B), MAD1L1, and MAD2L1 using a BioID2 proximity labeling proteomic approach (18) (Fig. 1*B*-*F*). The over-expression of critical cell division proteins often leads to cell division defects that can preclude the generation of epitope-tagged stable cell lines. Therefore, we first sought to generate BioID2 Gateway-compatible vectors with a doxycycline (Dox) inducible expression functionality. To do this, we amplified BirA-Myc with linkers coding for 27 or 47 amino acid residues downstream of Myc (BirA-Myc-27/47) (supplemental Fig. S1*A* and supplemental Table S2). These amplification products were cloned into the pGLAP1 vector (22), which had been previously modified by removal of its LAP-tag (EGFP-Tev-S-protein), to generate the pGBioID2-27 and pGBioID2-47 vectors (supplemental Fig. S1*A*). Full-length human open reading frames encoding for BUB1, BUB3, BUBR1, MAD1L1, and MAD2L1 were cloned into the pGBioID2-47 vector. The pGBioID2-47-SAC protein vectors (supplemental Fig. S1*B* and supplemental Table S3), were co-transfected with a vector expressing the Flp recombinase (pOG44) into HeLa Flp-In T-REx cells (supplemental Fig. S1*C*). Hygromycin resistant clones were then selected (supplemental Fig. S1*D*) and grown in the presence or absence of Dox for 16 hours. The Dox-induced expression of each BioID2-47-SAC protein was then assessed by immunoblot analysis (Fig. 2*A*). All of the BioID2-tagged core SAC proteins were expressed only in the presence Dox (Fig. 2*A*), indicating the successful establishment of inducible BioID2-tagged core SAC protein stable cell lines.

**Fig. 2.**
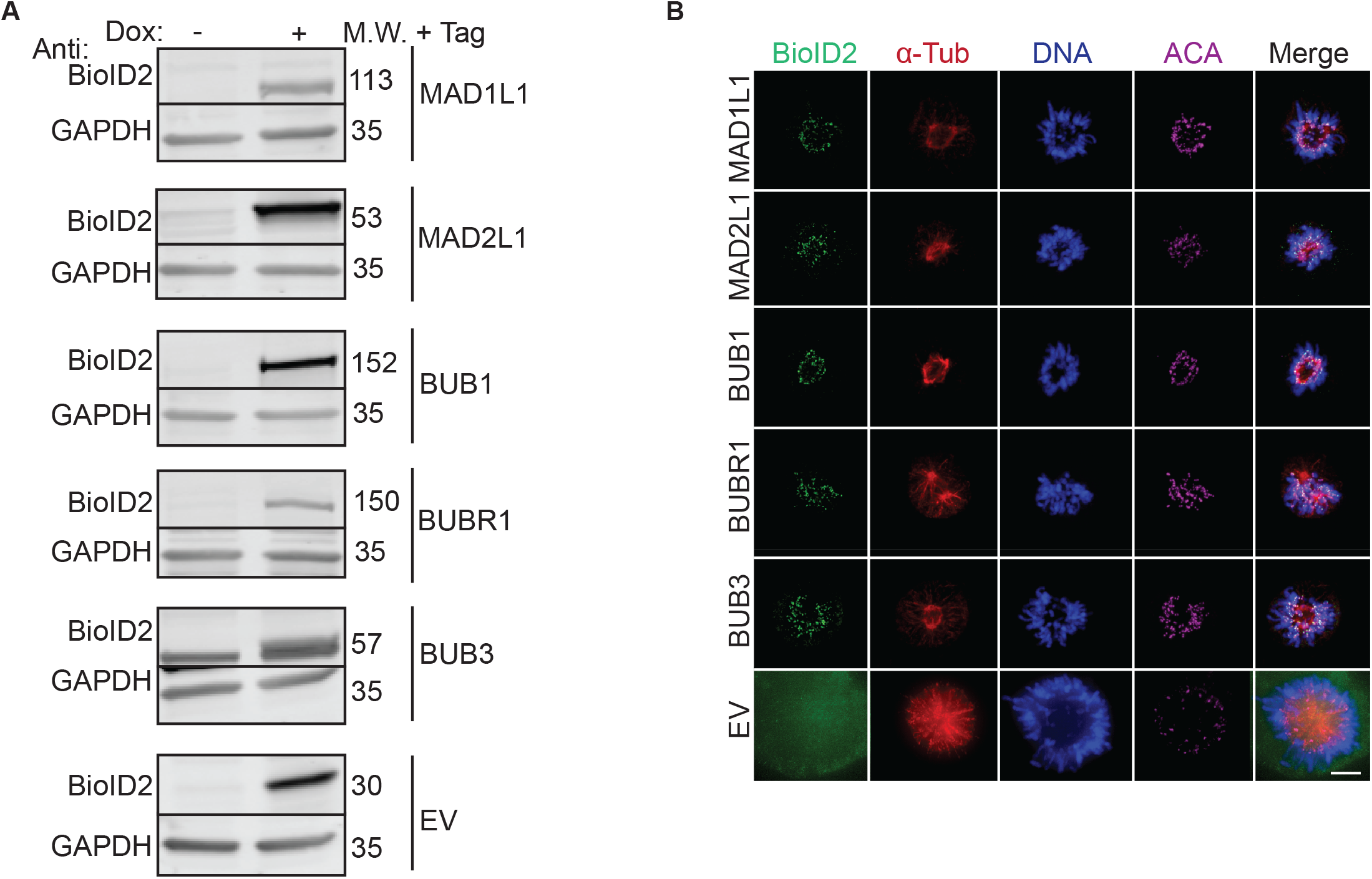
Establishment of inducible BioID2-tagged SAC protein (BUB1, BUB3, BUBR1, MAD1L1 and MAD2L1) stable cell lines. *A,* Immunoblot analysis of extracts from doxycycline (Dox)-inducible BioID2-tage alone or BioID2-tagged SAC protein (BUB1; BUB3; BUBR1; MAD1L1; MAD2L1) expression cell lines in the absence (−) or presence (+) of Dox for 16 hours. For each cell line, blots were probed with anti-BioID2 (to visualize the indicated BioID2-tagged SAC protein) and anti-GAPDH as a loading control. M.W. indicates molecular weight. Note that BioID2-tagged SAC proteins are only expressed in the presence of Dox. *B,* Fixed-cell immunofluorescence microscopy of the BioID2-tag alone or the indicated BioID2-tagged SAC proteins during prometaphase, a time when the SAC is active. HeLa BioID2-tagged protein expression cell lines were induced with Dox for 16 hours, fixed and stained with Hoechst 33342 DNA dye and anti-BioID2, anti-α-Tubulin and anti-centromere antibodies (ACA). Bar indicates 5*μ*m. Note that all BioID2-tagged SAC proteins localize to the kinetochore region (overlapping with the ACA signal), whereas the BioID2-tag alone was absent from kinetochores.

### BioID2-SAC Proteins Localize Properly to Kinetochores During Prometaphase

Next the ability of BioID2-SAC proteins to properly localize to the kinetochores during prometaphase, a time when the SAC is active and core SAC proteins localize to the kinetochore region, was analyzed by immunofluorescence microscopy. BioID2-SAC protein HeLa inducible stable cells lines were treated with Dox for 16 hours, fixed, and stained with Hoechst 33342 DNA dye and anti-BioID2, anti-α-Tubulin and anti-centromere antibodies (ACA). The localization of BioID2-SAC proteins in prometaphase cells was then monitored by immunofluorescence microscopy. BioID2-tagged BUB1, BUB3, BUBR1, MAD1L1, and MAD2L1 localized to kinetochores, overlapping fluorescence signal with anti-centromere antibodies (ACA) during prometaphase (Fig. 2*B*). In contrast, the BioID2-tag alone showed no specific localization (Fig. 2*B*). These results indicated that the BioID2-tag was not perturbing the ability of the SAC proteins to localize to kinetochores during the time when the SAC was active.

### BioID2-SAC Protein Proximity Labeling, Purifications, and Peptide Identification

To define the protein proximity networks of core SAC proteins, the inducible BioID2-SAC protein HeLa stable cell lines were used to perform BioID2-dependent proximity biotin labeling and biotinylated proteins were purified with a streptavidin resin (Fig. 3*A* and 3*B*). Briefly, inducible BioID2-SAC protein HeLa stable cells lines were treated with 0.2 μg/ml Dox, 100 nM Taxol, and 50 mM Biotin for 16 hours to induce the expression of BioID2-SAC proteins and to activate the SAC and arrest cells in prometaphase. Mitotic cells were collected, lysed, and the cleared lysates were bound to streptavidin beads. Bound biotinylated proteins were trypsinized on the beads and the peptides were analyzed by 2D-LC MS/MS (for details see Experimental Procedures). A diagnostic immunoblot analysis of each purification, using anti-BioID2 antibodies, showed that BioID2-tagged BUB1, BUB3, BUBR1, MAD1L1, and MAD2L1 were present in the extracts and were purified with the streptavidin beads, indicating that they had been biotinylated (Fig. 3*B*). In-house R scripts were then used to analyze the mass spectrometry results (for details see Experimental Procedures), to draw significance between peptides shared between the experimental and the control, we estimated the distribution of the mean difference of emPAI scores across proteins (for details see Experimental Procedures). Proteins that showed significant higher values in test purifications compared to the controls (values that lied outside of 95% confidence interval of the population mean difference) were considered hits and further analyzed (supplemental Table S4).

**Fig. 3.**
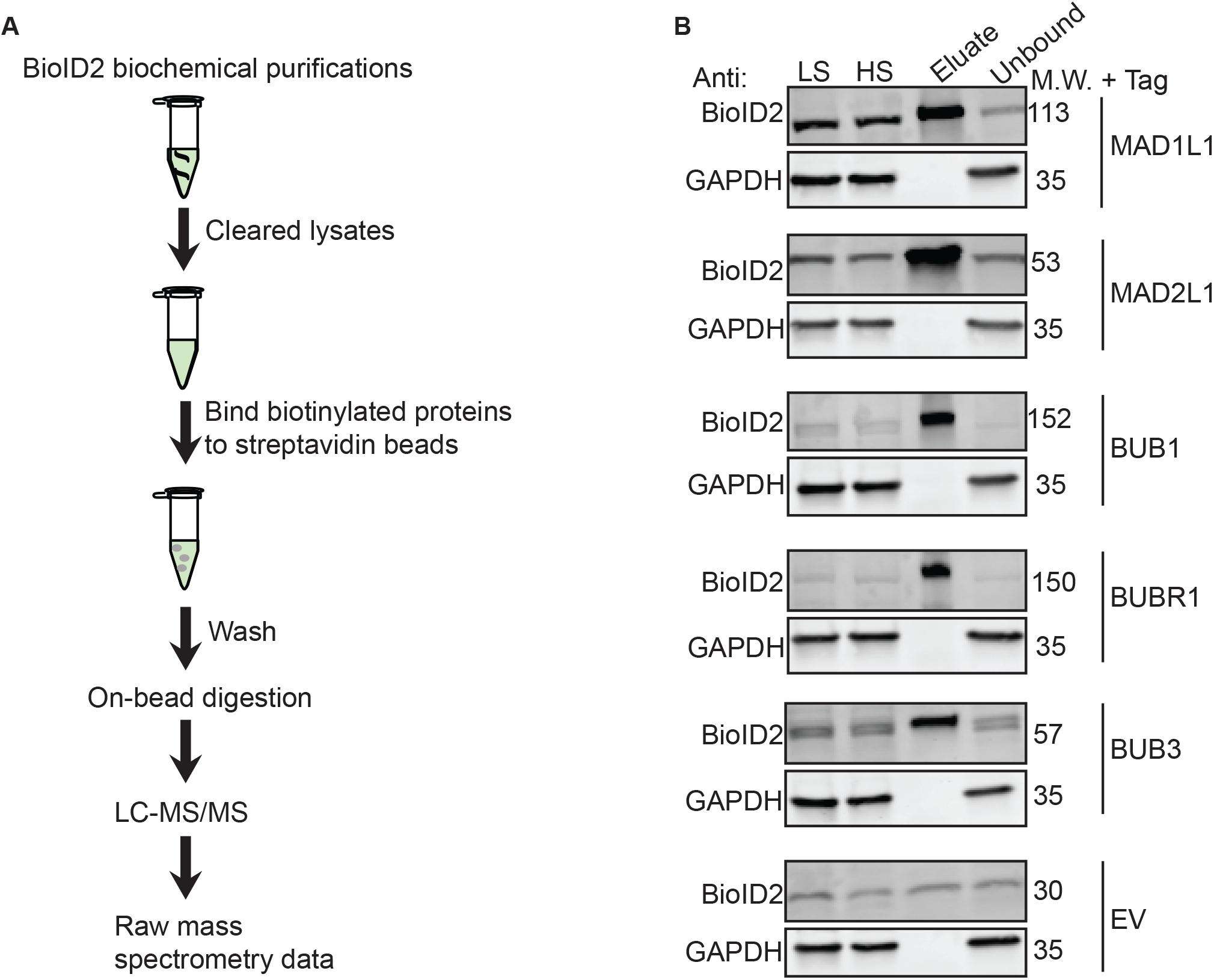
BioID2 biochemical purifications. *A,* HeLa BioID2-tag alone or BioID2-tagged SAC protein (BUB1; BUB3; BUBR1; MAD1L1; MAD2L1) stable cell lines were induced to express the BioID2-tagged proteins for 16 hours in the presence of 100 nM Taxol. BioID2 biochemical purifications were then performed with streptavidin beads to capture biotinylated proteins and the identification of biotinylated proteins was carried out by liquid chromatography-tandem mass spectrometry (LC-MS/MS) analysis. *B,* Immunoblot analysis of BioID2 biochemical purifications from cells expressing the indicated BioID2-tagged SAC proteins or the BioID2-tag alone. For each cell line, blots were probed with anti-BioID2 (to visualize the indicated BioID2-tagged SAC protein) and anti-GAPDH as a loading control. M.W. indicates molecular weight. LS indicates low speed supernatant, HS indicates high speed supernatant.

### Analysis of the Core SAC Protein Proximity Association Network

In-house R scripts were then used to integrate the identified proteins from the mass spectrometry analysis with the data visualization application RCytoscapeJS (32) to generate protein proximity association maps for each of the core SAC proteins (BUB1; BUB3; BUBR1; MAD1L1; MAD2L1) (supplemental Fig. S2). These five maps were compiled to generate the SAC proximity protein network (supplemental Fig. S3). To begin to digest the wealth of information within the SAC proximity protein network, we first analyzed the network with the CORUM database (28) and examined the proximal associations between each of the core SAC proteins. This analysis revealed many of the previously characterized core SAC component protein-protein interactions and the BUB1-BUB3, BUBR1-BUB3, BUBR1-BUB3-CDC20 (BBC subcomplex of the MCC) and MAD2L1-BUBR1-BUB3-CDC20 (MCC) complexes (Fig. 4 and supplemental Fig. S3) (6,36–38). These SAC complexes are critical to the establishment and maintenance of the SAC (39) and their identification was an indication that our proximity-based labeling approach was robust. Of interest, BUB3 was present in all of the purifications, consistent with its central role in recruiting other SAC proteins to the kinetochore and coordinating the formation of SAC sub-complexes (Fig. 4) (12). Although MAD1L1 and MAD2L1 had been previously determined to bind directly (40), our approach was unable to detect this association. However, previous proteomic analyses with N- or C-terminal BioID-tagged MAD1L1 were also unable to detect an association with MAD2L1, which was attributed to a low number of lysines in MAD2L1 that likely affected its efficient trypsin digestion (41).

**Fig. 4.**
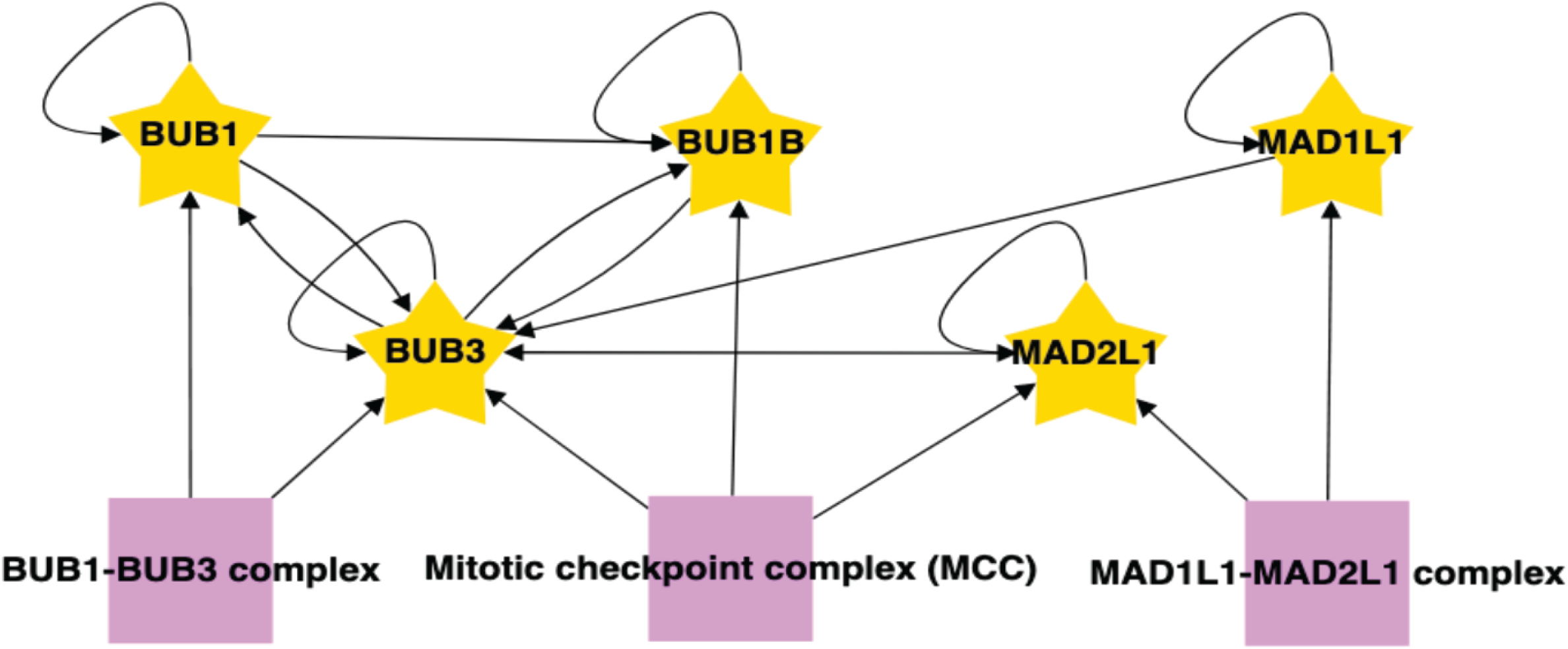
Associations among the core SAC proteins identified in the proximity protein network. The associations between each of the core SAC proteins (BUB1; BUB3; BUBR1; MAD1L1; MAD2L1) were isolated from the unified core SAC protein proximity association network (supplemental Fig. S3). Purple boxes highlight protein complexes known to assembly with core SAC proteins as annotated by the CORUM database. Arrows indicate the direction of the detected associations.

### Analysis of Core SAC Protein-Kinetochore Protein Proximity Associations

To specifically analyze the kinetochore proteins identified in the core SAC protein proximity networks, we applied a kinetochore related Gene Ontology (GO) annotation analysis to the data sets. Briefly, R scripts were used to integrate the identified proteins with the bioinformatic databases CORUM (28), Gene Ontology (30), BioGRID (29), and Reactome (31) using kinetochore related GO terms (see supplemental Table S5 for a list of Kinetochore GO IDs) to reveal the kinetochore associated proteins. RCytoscapeJS (32) was then used to generate GO, BioGRID, and Reactome kinetochore protein proximity association maps for each of the core SAC proteins (BUB1; BUB3; BUBR1; MAD1L1; MAD2L1) (supplemental Fig. S4-S8). The five kinetochore GO maps (one for each core SAC protein) were compiled to generate one core SAC protein kinetochore GO network that visualized the proteins within the network that were active at the kinetochore (supplemental Fig. S9*A*). A similar process was repeated to generate one core SAC protein BioGRID network that displayed the verified associations between the proteins that were active at the kinetochore (supplemental Fig. S9*B*) and one core SAC protein Reactome network that highlighted the cellular pathways that proteins in the SAC proximity association network have been linked to (supplemental Fig. S9*C*). Additionally, we generated core SAC protein GO, BioGRID, and Reactome networks using mitotic spindle related GO annotations (see supplemental Table S5 for a list of mitotic spindle GO IDs) (supplemental Fig. S10*A-C*). Finally, we generated core SAC protein GO, BioGRID, and Reactome networks using both the kinetochore and mitotic spindle related GO annotations (Fig. 5*A-C*). Together, these networks not only visualized the associations of each core SAC protein with kinetochore components and more broadly proteins implicated in mitotic spindle assembly, they also provided a holistic view of their interconnectedness (ie. associations among core SAC proteins and subcomplex and complex formation).

**Fig. 5.**
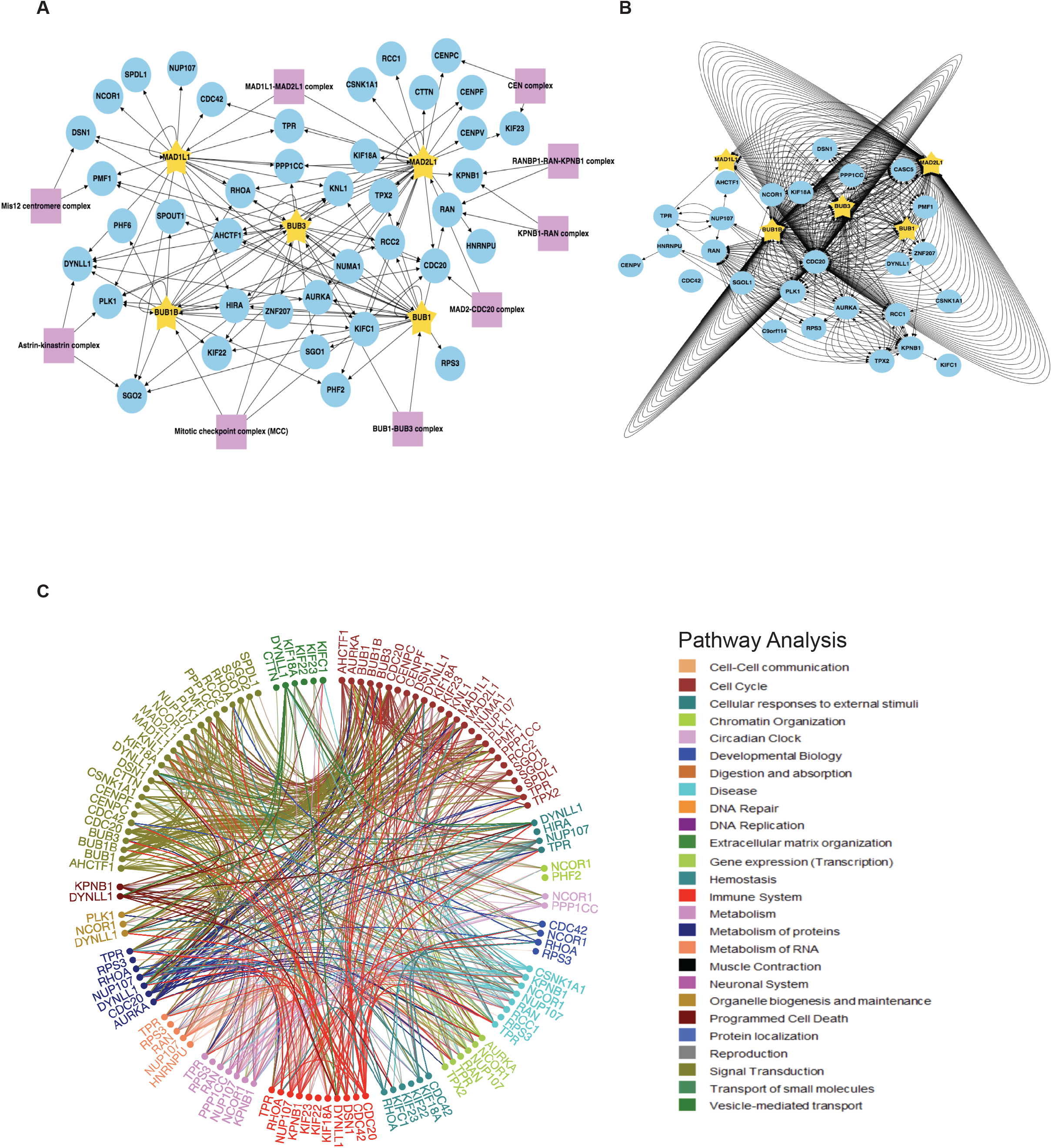
SAC protein BioID2 kinetochore/mitotic spindle assembly proximity association network. *A,* Individual core SAC protein (BUB1; BUB3; BUBR1; MAD1L1; MAD2L1) proximity protein maps were compiled and subjected to kinetochore and mitotic spindle assembly GO annotation analysis along with a COURM complex annotation analysis to generate a core SAC protein kinetochore/mitotic spindle assembly proximity association network. Purple boxes highlight kinetochore and/or mitotic spindle assembly associated protein complexes present in the network. Arrows indicate the direction of the detected interactions. For a list of GO terms used see supplemental Table S5. *B,* The core SAC protein kinetochore/mitotic spindle assembly proximity association network was analyzed with BioGRID to reveal previously verified protein associations. Each arrow indicates an experimentally annotated interaction curated in the BioGRID database. Direction of arrows indicate an annotated interaction from a bait protein to the prey. *C,* Reactome pathway analysis of the core SAC protein kinetochore/mitotic spindle assembly proximity association network. The Reactome circular interaction plot depicts the associations between the identified proteins within the SAC protein kinetochore/mitotic spindle assembly proximity association network and the corresponding pathways in which they function. Legend presents the color-coded pathways that correspond to the circular interaction plots.

Numerous insights were derived from these networks and we highlight four here. First, we identified the Mis12 centromere complex components DSN1 and PMF1 in the BUB1 and MAD1L1 purifications (Fig. 5*A* and supplemental Figs. S4*A* and S7*A*). The Mis12 complex is comprised of PMF1, MIS12, DSN1, and NSL1 (42–44) and genetic and biochemical studies have shown that it coordinates communication from the outer kinetochore to the centromeric DNA in the inner kinetochore (44–46). Unexpectedly, PMF1 was also identified in the BUB3 purification (Fig. 5*A* and supplemental Fig. S5*A*). To our knowledge there have been no previous reports of a direct association between BUB3 and the Mis12 complex. Therefore, this BUB3-PMF1 association could indicate a novel direct interaction or simply that these proteins reside within close proximity at the kinetochore. Of interest, the Mis12 complex recruits KNL1 to the kinetochore, which functions as a scaffold for the recruitment of BUB3 that subsequently recruits additional SAC components (4,38,47). Consistently, we observed the association of KNL1 with BUB1, BUB3, BUBR1, and MAD1L1 (Fig. 5*A*). These associations were previously reported, as summarized in the Figure 5*B* BioGRID network, and had been established to have a role in checkpoint activation (41,48–50) (reviewed in (5)). Additionally, MAD2L1 was not found to associate with KNL1 and to our knowledge a KNL1-MAD2L1 interaction has not been reported.

Second, minor components of the Astrin-Kinastrin complex (PLK1, DYNLL1, and SGO2) (51) were found to associate with all of the core SAC proteins (Fig. 5*A* and supplemental Figs. S4*A,* S5*A,* S6*A,* S7*A,* S8*A*) and more significantly with BUB1 (supplemental Fig. S11). The Astrin-Kinastrin complex is important for aligning and attaching microtubules to kinetochores (51–53). Previous studies showed that depletion of BUB1 led to the delocalization of PLK1 and SGO2 from the kinetochores during prometaphase (54,55). Additionally, the BUB1 kinase activity was shown to be important for SGO2 kinetochore localization (56) and for the proper localization of BUB1 to the kinetochore (55) and pharmacological inhibition of the BUB1 kinase activity led to delocalization of SGO2 away from kinetochores (57). However, whether the BUB1 kinase activity was required for PLK1 kinetochore localization remained unknown. To address this, we first sought to confirm that PLK1 and SGO2 were mislocalized in BUB1-depleted cells. Indeed, immunofluorescence microscopy of HeLa cells treated with control siRNA (siControl) or BUB1-targeting siRNA (siBUB1) showed that both PLK1 and SGO2 were absent from kinetochores in siBUB1-treated prometaphase cells (Fig. 6*A* and 6*B*). Next, we asked if the BUB1 kinase activity was required for PLK1 and SGO2 kinetochore localization. RPE cells were treated with control DMSO vehicle or the recently developed BUB1 kinase selective inhibitor BAY 1816032 (21) and the localization of PLK1 and SGO2 was assessed in mitotic cells. In comparison to control DMSO-treated cells, PLK1 and SGO2 were absent from kinetochores in BAY 1816032-treated cells (Fig. 6*C* and 6*D*). Additionally, when BioID2-BUB1 expressing HeLa cells were treated with BAY 1816032, the BioID2-BUB1 kinetochore signal was decreased (Fig. 6*E*). This data indicated that the BUB1 kinase activity was important for the its proper localization to kinetochores and for the localization of the Astrin-Kinastrin minor complex components PLK1 and SGO2 to the kinetochore.

**Fig. 6.**
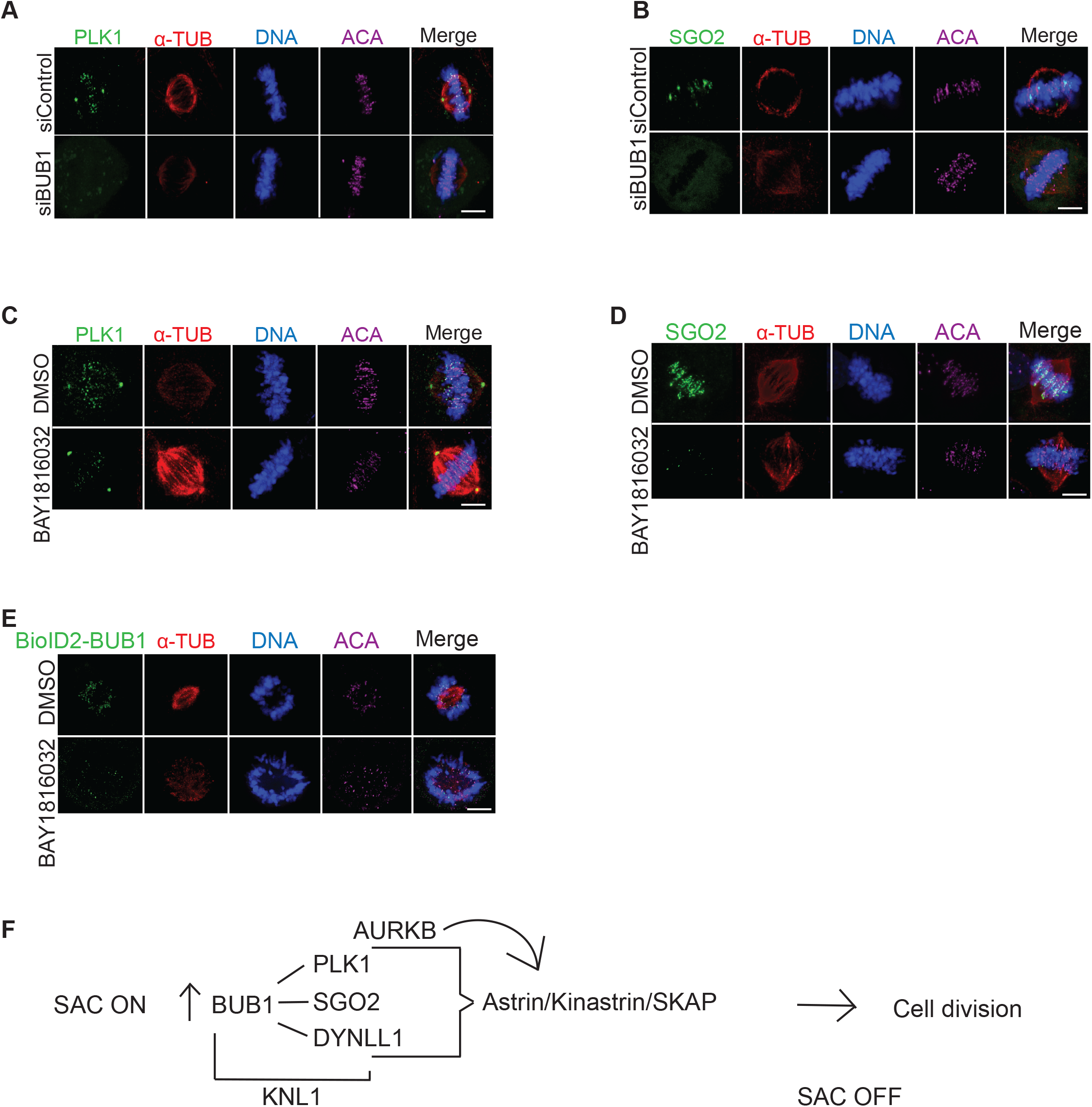
BUB1 as a hub for organizing the metaphase to anaphase transition. *A-B*, Fixed-cell immunofluorescence microscopy of mitotic HeLa cells treated with control siRNA (siControl) or siRNA targeting BUB1 (siBUB1). Cells were fixed and stained with Hoechst 33342 DNA dye and anti-PLK1 (*A*) or SGO2 (*B*) antibodies, along with anti-α-Tubulin and anti-centromere antibodies (ACA). Bars indicate 5*μ*m. Note that PLK1 and SGO2 are delocalized from the kinetochore region (overlapping with the ACA signal) in siBUB1-treated cells. *C-D*, Same as in *A*, except that RPE cells were used and treated with control DMSO vehicle (*C*) or the BUB1 kinase inhibitor BAY 1816032 (*D*). Note that PLK1 and SGO2 are delocalized from the kinetochore region in BAY 1816032-treated cells. Bars indicate 5*μ*m. *E*, Same as in C, except that a HeLa BioID2-BUB1 expressing cell line was used. Bar indicates 5*μ*m *F*, Model of BUB1 as an organizer of the metaphase to anaphase transition. BUB1 is critical for SAC protein binding to KNL1 to establish the SAC response and is also critical for the recruitment of the Astrin-Kinastrin minor complex, which is essential for the metaphase to anaphase transition.

Third, we identified CENPV as a MAD2L1 associating protein (Fig. 5*A*). CENPV was identified in a proteomic screen for novel components of mitotic chromosomes (58) and was later shown to localize to kinetochores early in mitosis and to have a major role in directing the chromosomal passenger complex (CPC) subunits Aurora B and INCENP to the kinetochore (50,59). Although BUB1 has been shown to be important for the recruitment of the CPC to kinetochore (60), we are unaware of any reports of MAD2L1 being involved in this process. Interestingly, MAD2L1 has been shown to regulate the relocation of the CPC from centromeres through its inhibition of MKLP2, which is essential for proper cytokinesis (61). Thus, it is possible MAD2L1 could also be regulating CPC localization to kinetochores through its association with CENP-V.

Fourth, components of the nuclear pore complex were found to associate with MAD1L1 and MAD2L1 (supplemental Fig. S12). To better visualize these nuclear pore associated proteins, we performed a proximity protein mapping analysis for each of the core SAC proteins using the nuclear pore related GO annotations (see supplemental Table S5 for a list of nuclear pore related GO IDs) (supplemental Fig. S12). This analysis revealed that MAD1L1 had associations with nuclear pore basket components including TPR, NUP153, NUP50, and other components of the nuclear pore that are in close proximity to the nuclear basket like ELYS/AHCTF1 (also known as MEL-28 in *C. elegans*) and NUP107 (supplemental Fig. S12). These data support previous studies in humans and other organisms that have shown that MAD1L1 associates with TPR, NUP153, and NUP107 and is important for generating the MAD1L1-MAD2L1 complex in early mitosis to establish the SAC (62–68). Similarly, MAD2L1 was found to associate with TPR (previously verified in (63), NUP50, Nup153, NUP210 and ELYS (supplemental Fig. S12). Of interest, we did not detect associations between other core SAC proteins (BUB1; BUB3; BUBR1) and nuclear pore basket proteins, with the exception of ELYS. These data are consistent with a model where MAD1L1 makes multiple direct contacts with the nuclear pore basket complex subunits and MAD2L1 is in close proximity to NUP153 and NUP50 due to its binding to MAD1L1. We note that ELYS was found in all of the core SAC protein proximity maps (supplemental Fig. S12). ELYS was discovered in a proteomic screen for NUP107-160 complex binding partners and was shown to localize to nuclear pores in the nuclear lamina during interphase and to kinetochores during early mitosis, similar to the NUP107-160 complex (69). More recently, ELYS was shown to function as a scaffold for the recruitment of Protein Phosphatase 1 (PP1) to the kinetochore during M-phase exit, which was required for proper cell division (70,71). However, it remains unclear what the connection is between ELYS and the core SAC proteins and whether ELYS plays a role in establishing or silencing the SAC.

### Core SAC Proteins in Cellular Homeostasis

It’s important to note that most of the core SAC proteins have been shown to have roles in cellular homeostasis independent of their role in the SAC, which are predominantly mediated through protein-protein interactions with non-kinetochore proteins. Many of these associations were present in the individual core SAC protein proximity maps where GO annotations were not applied (supplemental Fig. S2). Consistently, Reactome pathway analysis of the core SAC protein proximity protein network showed that many of the SAC associated proteins had roles in numerous pathways important for cellular homeostasis including the cell cycle, DNA repair, and gene expression (Fig. 5*C*). Finally, we emphasize that researchers will be able to interrogate the core SAC protein proximity networks to uncover other potentially important associations that will inform on the function and regulation of the SAC. All mass spectrometry data files and R scripts have been deposited in open access web servers, for details see Experimental Procedures.

## DISCUSSION

The SAC is an important signaling pathway that is critical for proper cell division, which functions with great precision in a defined spatial and temporal manner (2). Due to the dynamic nature of the associations between core SAC proteins and the complexes and subcomplexes that they form, it has been difficult to generate a comprehensive proteomic view of the proteins that are in close proximity and that interact with core SAC proteins. Here, we have established an inducible BioID2-tagging system that allowed for the transient expression of BioID2-tagged core SAC proteins (BUB1, BUB3, BUBR1, MAD1L1, and MAD2L1), which bypasses issues associated with long-term overexpression of key cell division proteins that can compromise cellular homeostasis. We coupled this system to a proximity labeling proteomic approach to systematically define a proximity protein association map for each of the core SAC proteins. These proximity maps were integrated to generate a core SAC protein proximity protein network. The coupling of the proximity maps/network with curated functional databases like CORUM, GeneOntology, BioGRID, and Reactome allowed for a systems level bioinformatic analysis of the associations within these maps/network. To our knowledge this is the first systematic characterization of the core SAC proteins by proximity-based proteomics.

Our analysis recapitulated many of the core SAC protein-protein interactions, sub-complexes, and complexes that had been previously described. Importantly, it also identified numerous novel associations that warrant further examination. Among these is ELYS, which associated with all of the core SAC proteins. Although an interpretation of these associations could be that the core SAC proteins associate with ELYS at the nuclear pore in preparation for mitotic entry and SAC activation, we favor a model where ELYS may be important for the recruitment of core SAC proteins to the kinetochore and/or for checkpoint activation. Future studies aimed at addressing these models should bring clarity to the potential role of ELYS in SAC functioning and cell division. Of interest, previous studies had shown the importance of BUB1 for the localization of the Astrin-Kinastrin minor complex proteins to the kinetochore (51–54) and our analysis further determined that the BUB1 kinase activity was important for this function. Together, these data indicate that BUB1 may have a central organizing role not only in SAC activation and function, but in SAC silencing and mediating the transition from metaphase to anaphase through its association with the Astrin-Kinastrin minor complex (Fig. 6*F*).

To facilitate the use and interrogation of these core SAC protein proximity maps/network by the scientific community, all mass spectrometry data and R scripts used to analyze the data have been deposited in open access databases (see Experimental Procedures). These tools will enable researches to define novel associations and to generate testable hypotheses to further advance the current understanding of SAC function and regulation.

## Abbreviations

BioID2: Biotin identification 2
SAC: Spindle assembly checkpoint

## ACKNOWLEDGEMENTS

This material is based upon work supported by the National Institutes of Health NIGMS grant number R01GM117475 to J.Z.T., any opinions, findings, and conclusions or recommendations expressed in this material are those of the authors and do not necessarily reflect the views of the National Institutes of Health NIGMS. Y.A.G. and K.M.C. were supported by the UCLA Tumor Cell Biology Training Program (USHHS Ruth L. Kirschstein Institutional National Research Service Award # T32CA009056). This work was supported in part by a grant to The University of California, Los Angeles from the Howard Hughes Medical Institute through the James H. Gilliam Fellowships for Advanced Study Program (E.F.V), by a UCLA Molecular Biology Institute Whitcome Fellowship (E.F.V.) and a NIH P30 DK063491 grant (J.P.W.).

