## Supplemental Material for "Dissection of the Spindle Assembly Checkpoint by Proximity Proteomics"

### Supplemental Figures

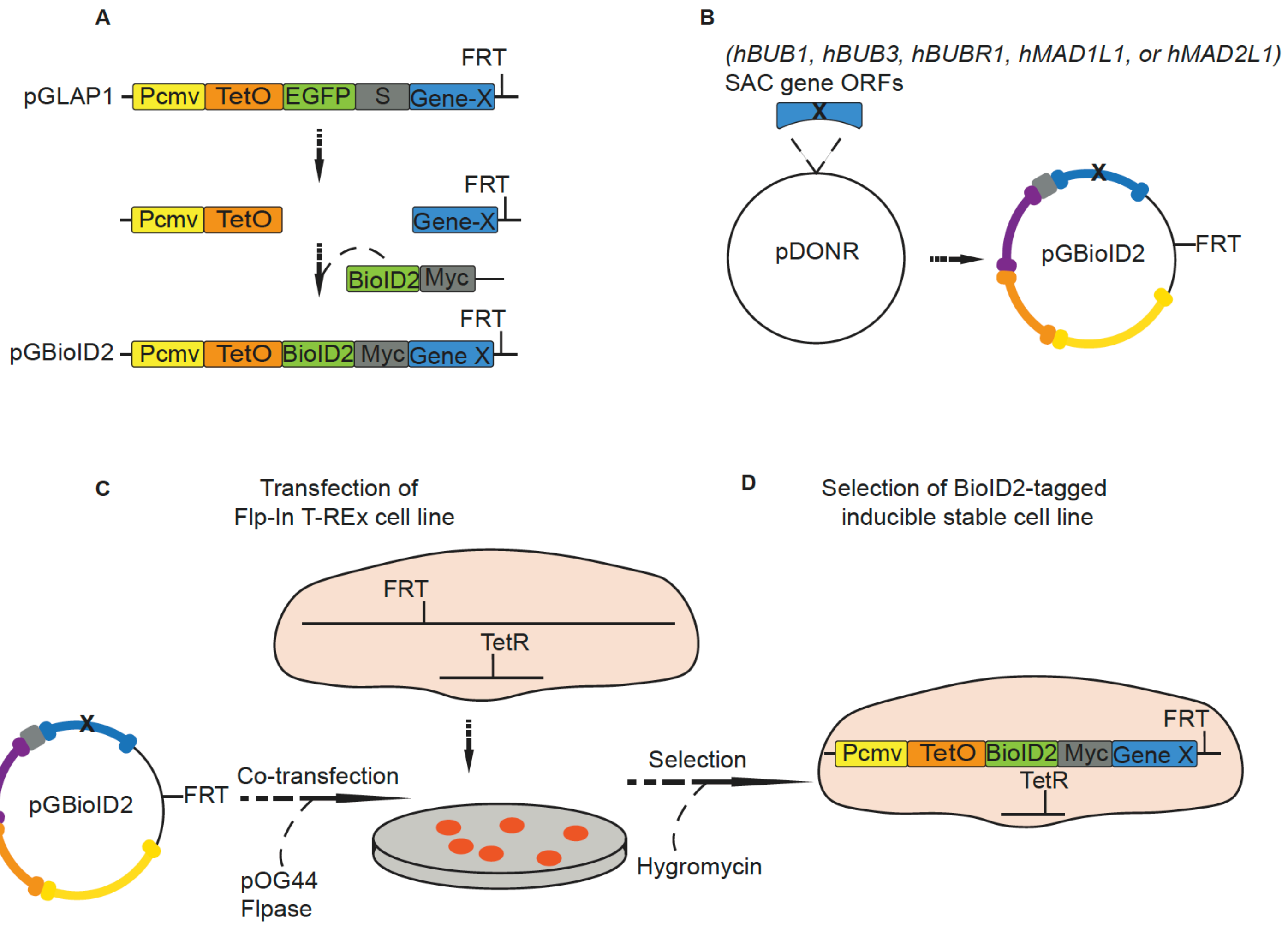

Supplemental Fig. S1. **Generation of BioID2-tagged SAC protein inducible stable cell lines.** *A,* Generation of the Gateway compatible pGBioID2 vectors (pGBioID2-27 and pGBioID2-47) using the pGLAP1 vector backbone. S denotes S-tag and M denotes Myc-tag. *B,* Transfer of core human SAC gene ORFs from the pDONR221 vector to the pGBioID2 vector. *C-D*, pGBioID2-SAC gene vectors were co-transfected with the Flpase containing vector pOG44 into HeLa Flp-In-TRex cells and stable integrants were selected with Hygromycin. See supplemental Table S2 and S3 for primer amplification sequences and information on the vectors generated.

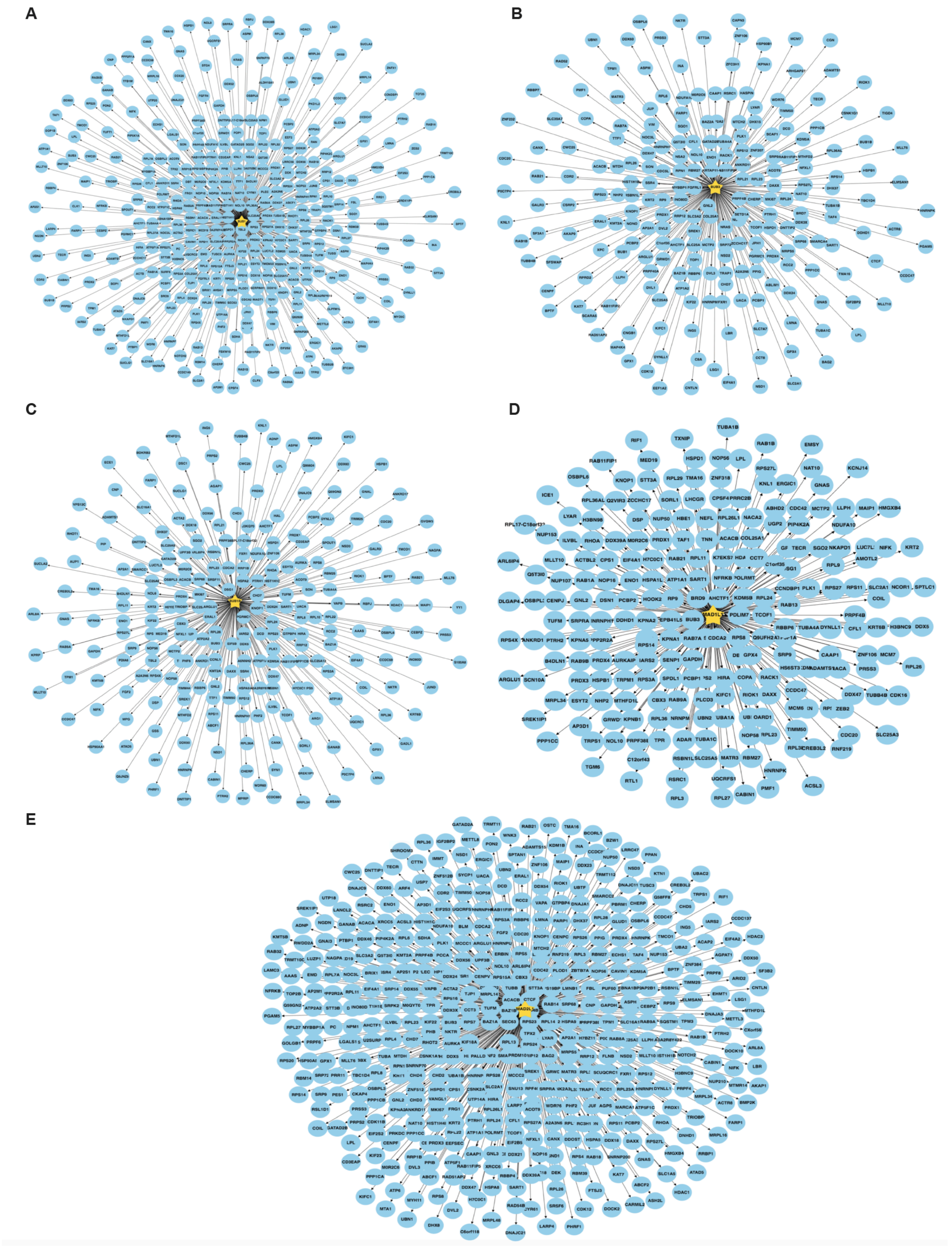

Supplemental Fig. S2. **Core SAC protein proximity association maps.** *A-E,* Protein proximity association maps for each of the core SAC proteins BUB1 (*A*), BUB3 (*B*), BUBR1 (*C*), MAD1L1 (*D*), and MAD2L1 (*E*). See supplemental Table S4 for a list of proteins in each map. The maps were visualized using RCytoscapeJS.

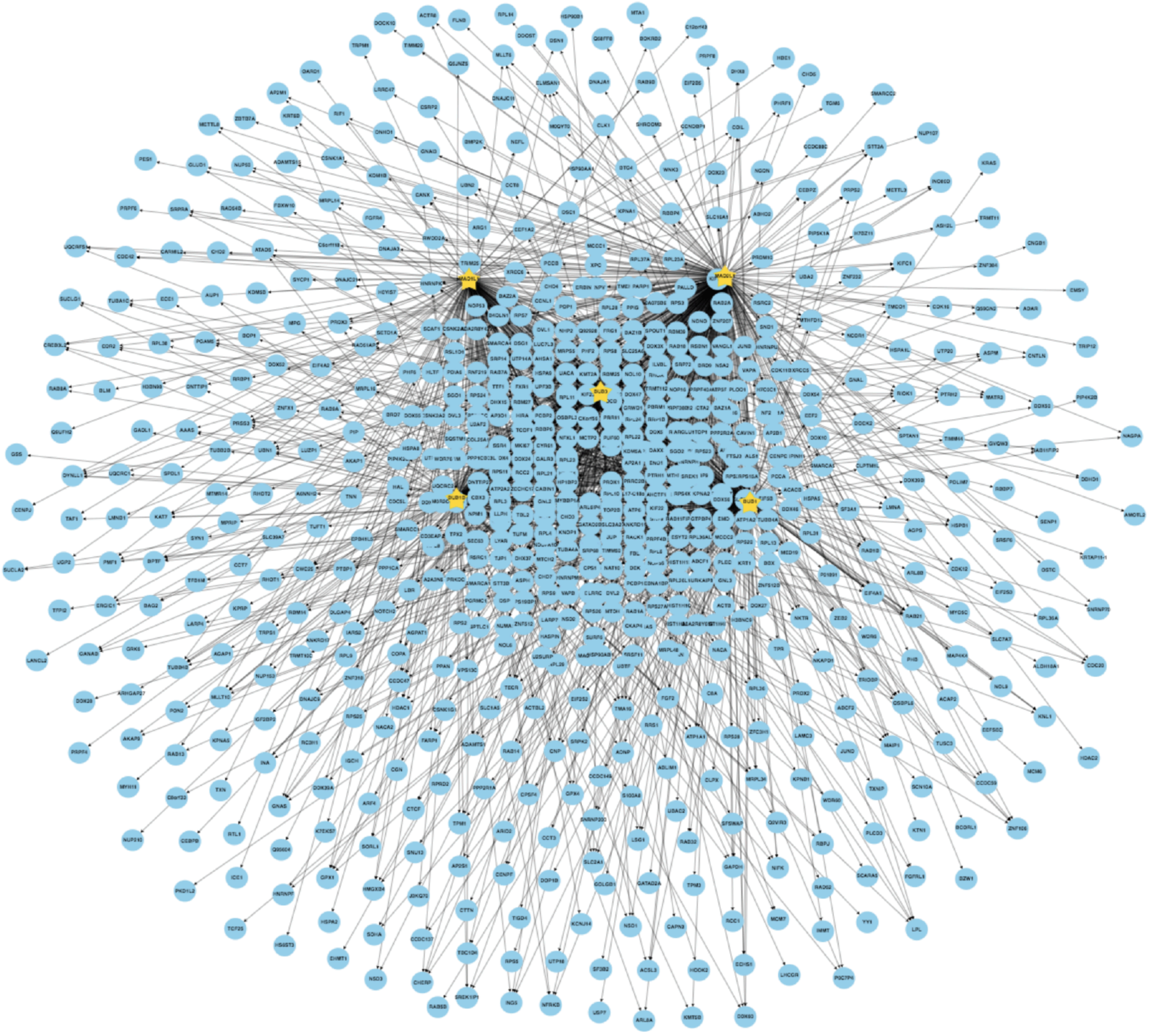

Supplemental Fig. S3. **Core SAC protein proximity network.** The five core SAC protein (BUB1; BUB3; BUBR1; MAD1L1; MAD2L1) proximity association maps in supplemental Figure S2 were combined to generate the core SAC protein proximity association network. The network was visualized using RCytoscapeJS.

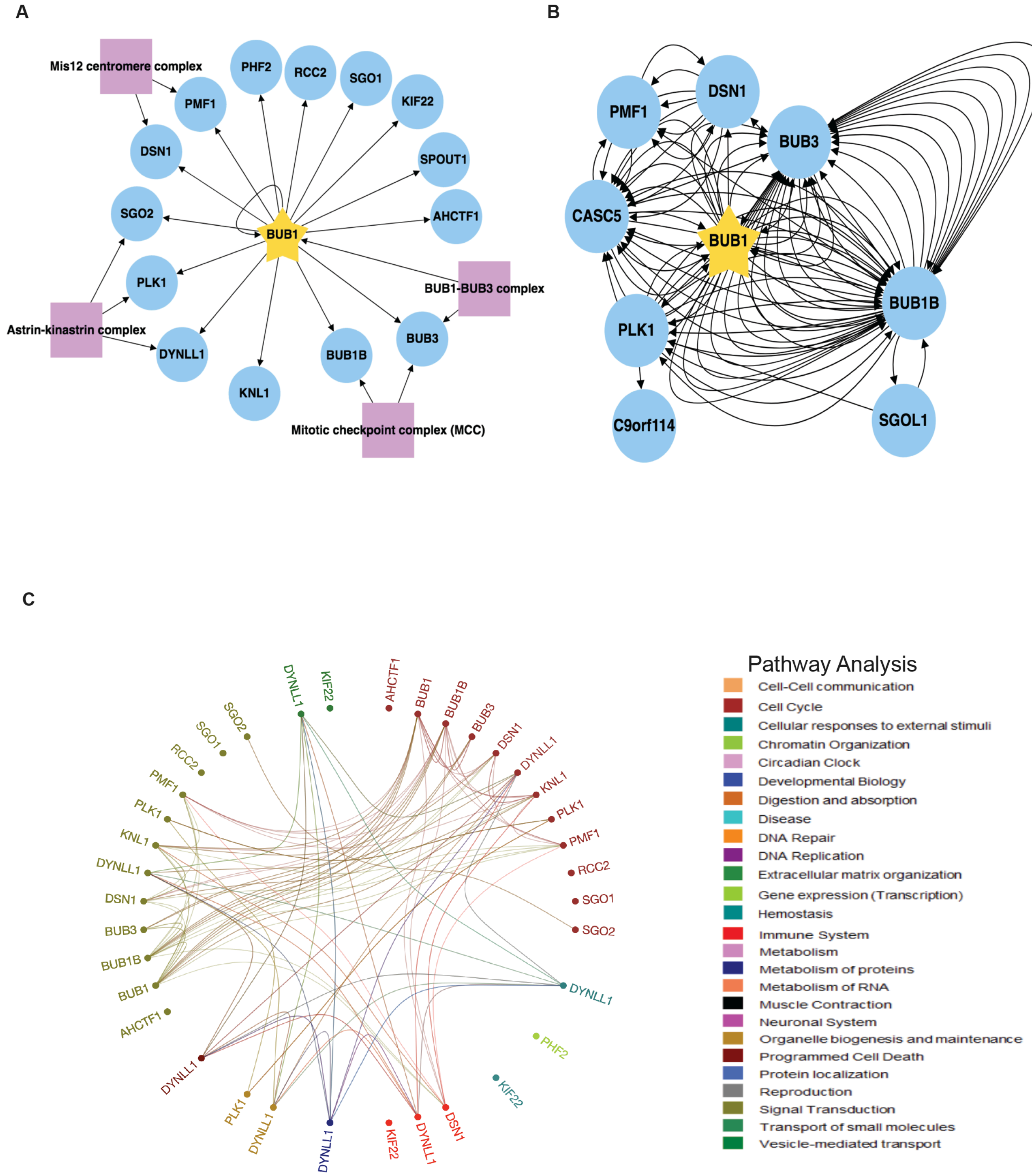

Supplemental Fig. S4. **The BioID2-BUB1 proximity protein association map.** *A*, Generation of the BUB1 protein proximity association map using kinetochore related Gene Ontology annotations and CORUM complex annotation analyses. The map was visualized using RCytoscapeJS. Purple boxes highlight protein complexes known to assemble with BUB1 as annotated by the CORUM database. Arrows indicate the direction of the detected interactions. *B,* The BUB1 proximity association map was analyzed with BioGRID to reveal previously verified protein associations. Each arrow indicates an experimentally annotated interaction curated in the BioGRID database. Direction of arrows indicate an annotated interaction from a bait protein to the prey. *C,* Reactome pathway analysis of the BUB1 proximity association map. The Reactome circular interaction plot depicts the associations between the identified proteins within the BUB1 proximity association map and the corresponding pathways in which they function. Legend presents color-coded pathways that correspond to the circular interaction plot.

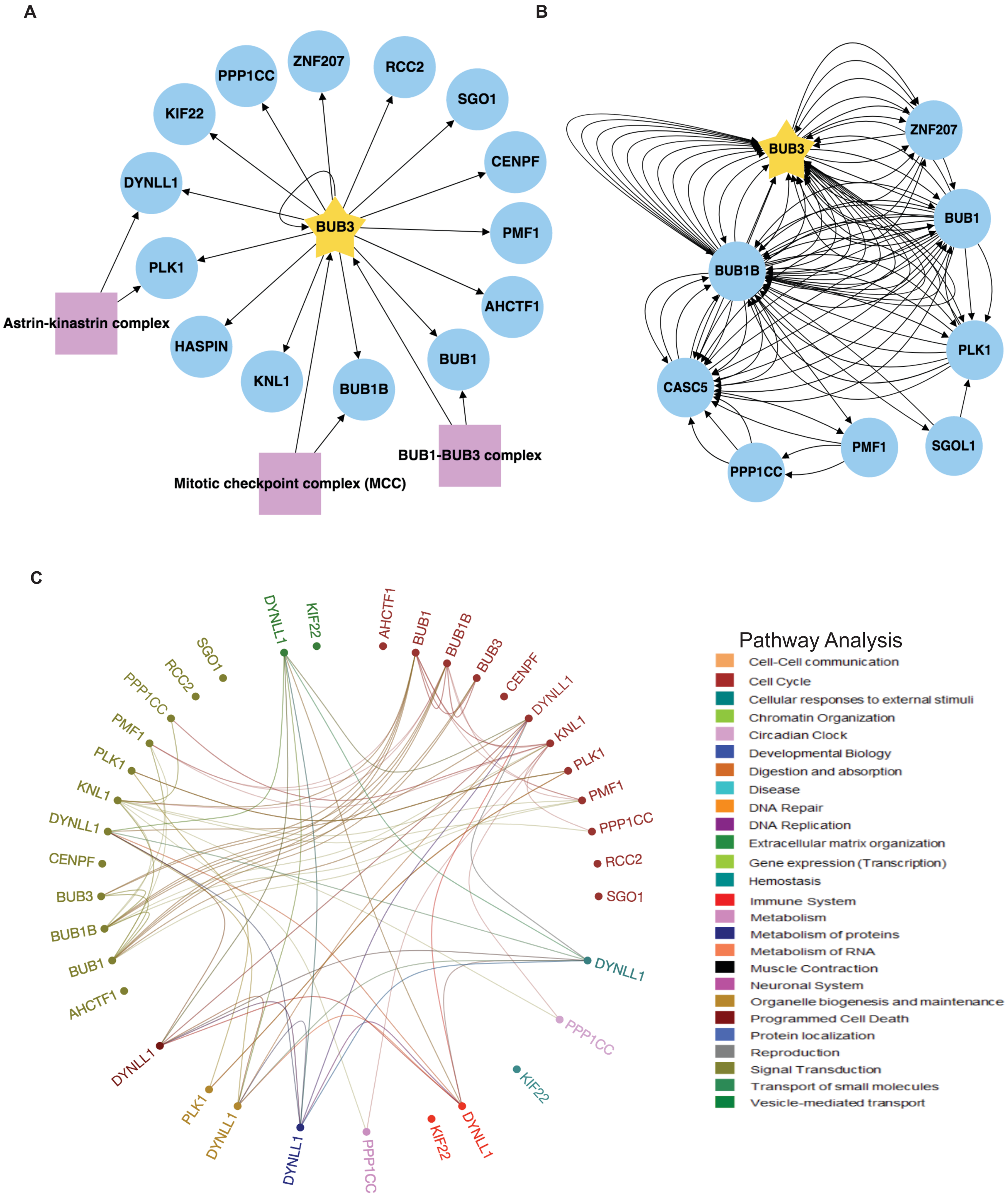

Supplemental Fig. S5. **The BioID2-BUB3 proximity protein association map.** *A*, *B*, and *C* are as described in supplemental Fig. S4.

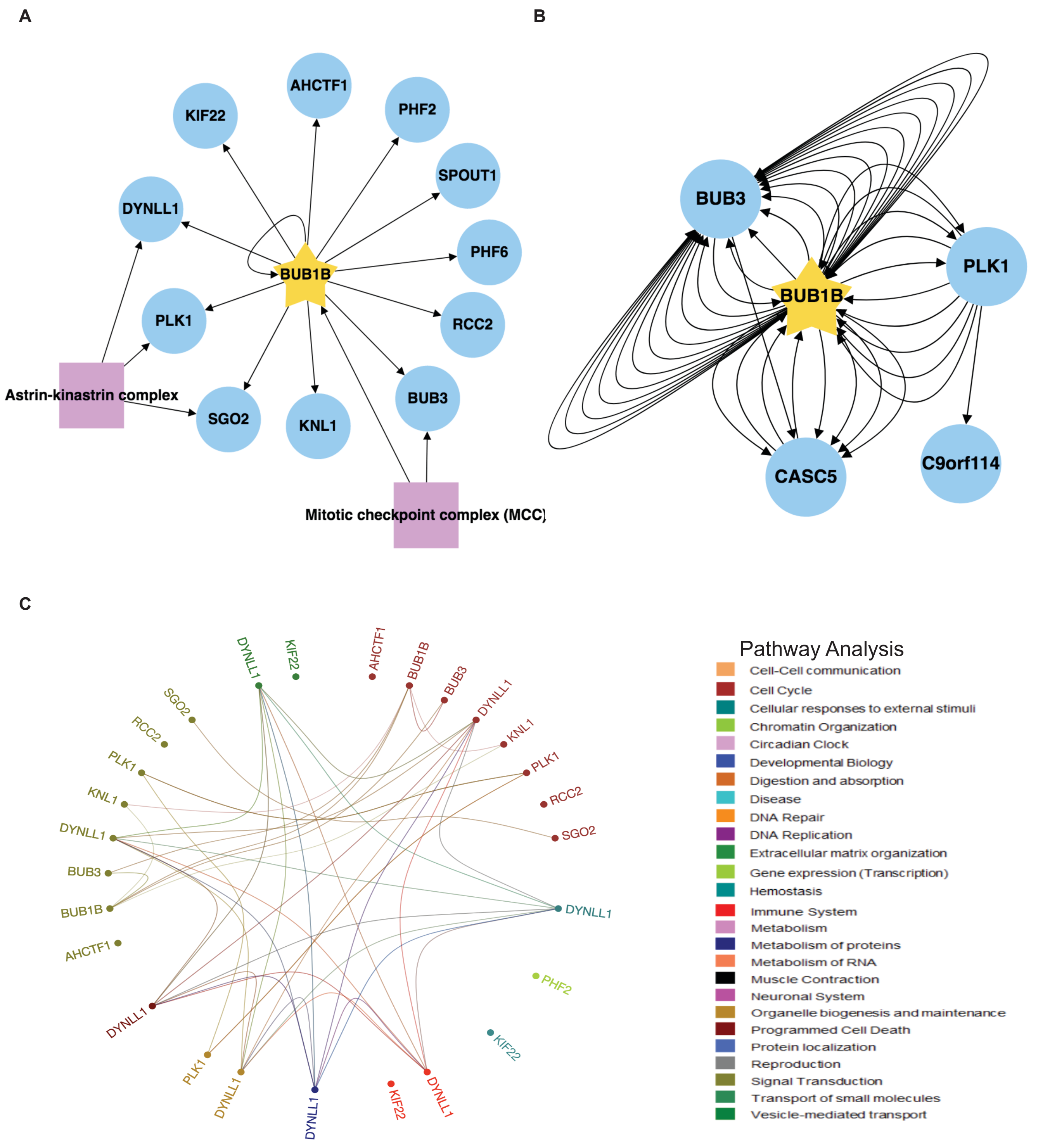

Supplemental Fig. S6. **The BioID2-BUBR1/BUB1B proximity protein association map.** *A*, *B*, and *C* are as described in supplemental Fig. S4.

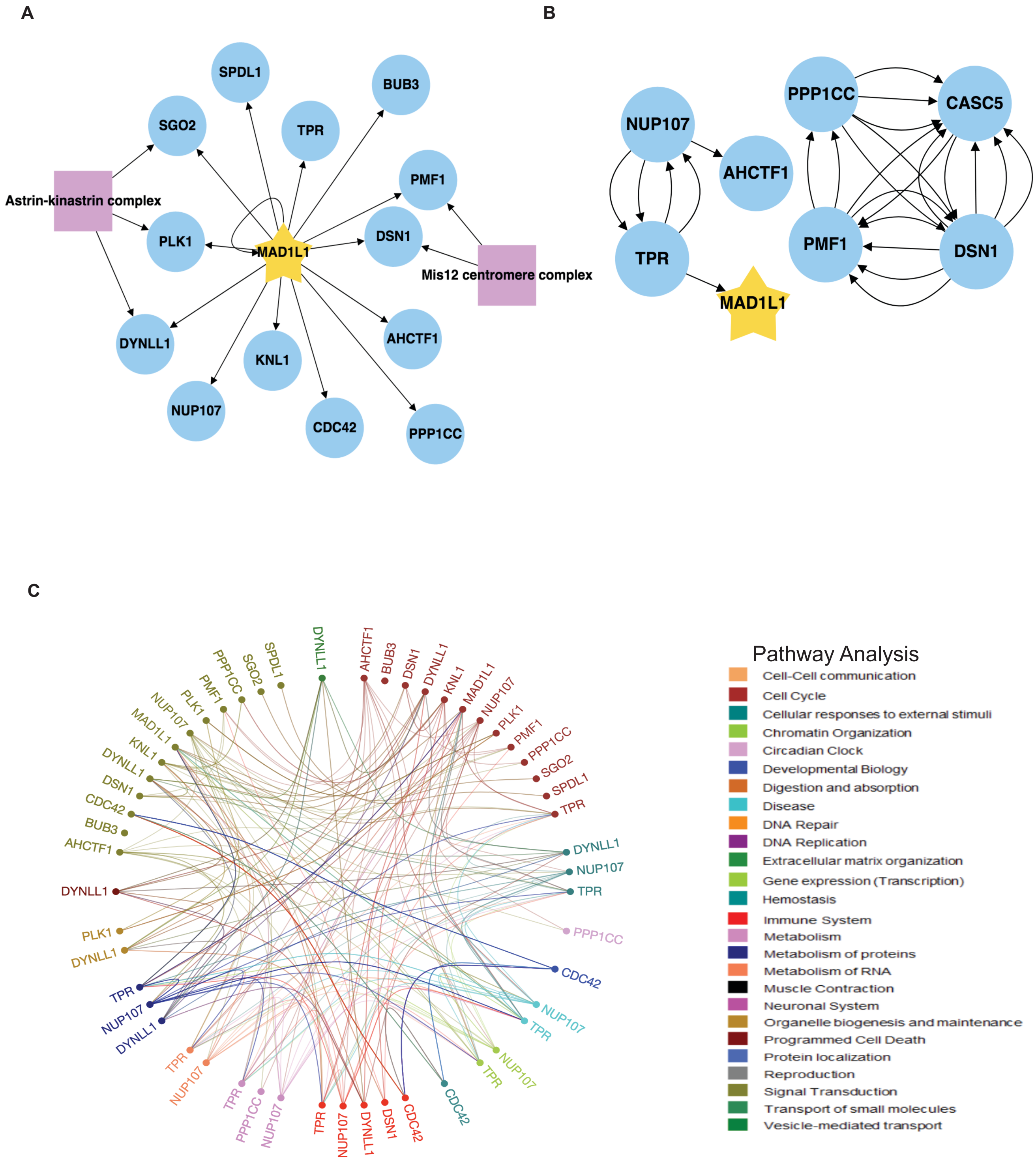

Supplemental Fig. S7. **The BioID2-MAD1L1 proximity protein association map.** *A*, *B*, and *C* are as described in supplemental Fig. S4.

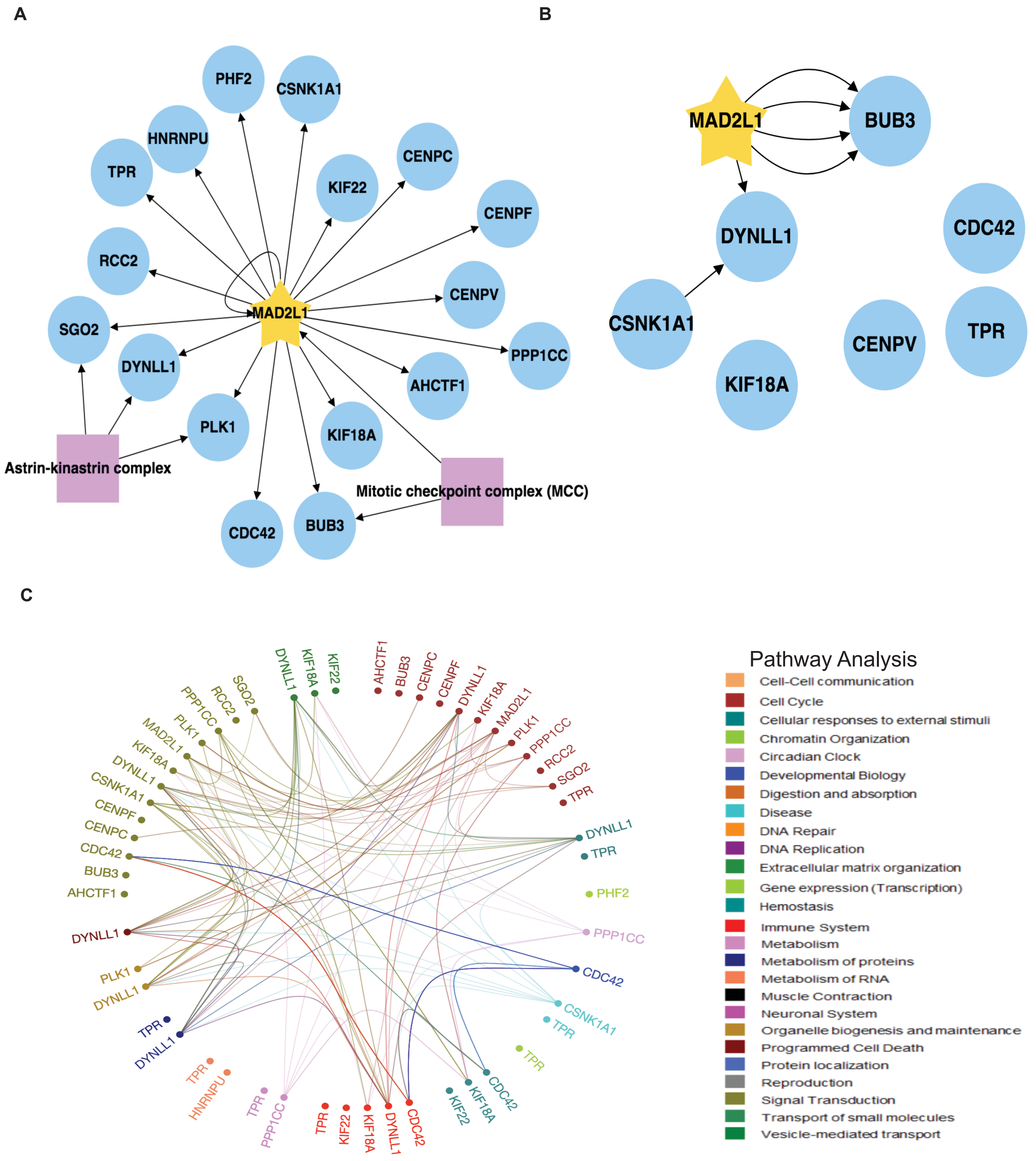

Supplemental Fig. S8. **The BioID2-MAD2L1 proximity protein association map.** *A*, *B*, and *C* are as described in supplemental Fig. S4.

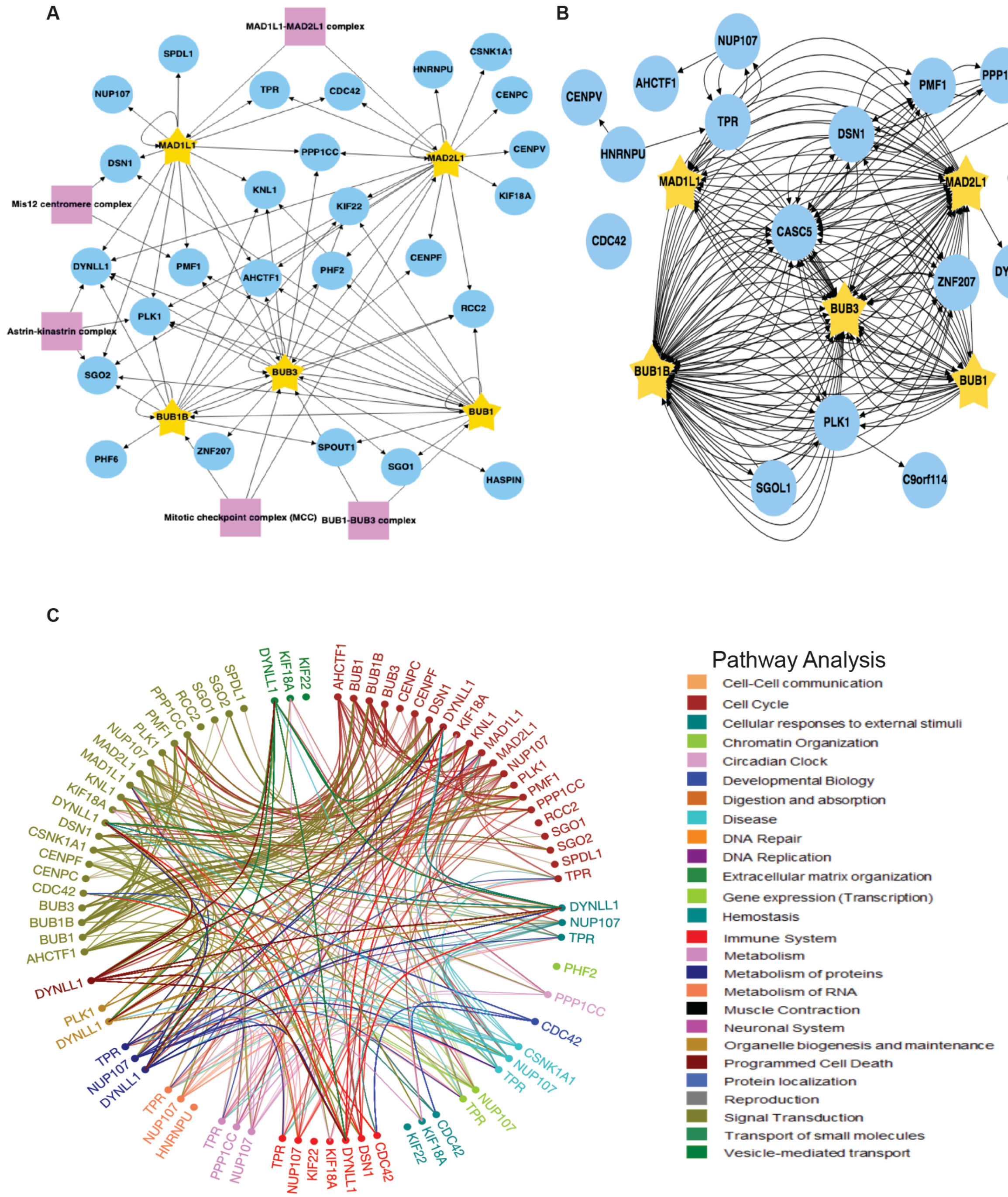

Supplemental Fig. S9. **The core SAC protein proximity association network using kinetochore Gene Ontology annotations.** *A*, Generation of the core SAC protein (BUB1; BUB3; BUBR1; MAD1L1; MAD2L1) proximity association network using kinetochore related Gene Ontology annotations and CORUM complex annotation analyses. The map was visualized using RCytoscapeJS. Purple boxes highlight protein complexes known to assembly with core SAC proteins as annotated by the CORUM database. Arrows indicate the direction of the detected interactions. *B,* The core SAC protein kinetochore proximity association network was analyzed with BioGRID to reveal previously verified protein associations. Each arrow indicates an experimentally annotated interaction curated in the BioGRID database. Direction of arrows indicate an annotated interaction from a bait protein to the prey. *C,* Reactome pathway analysis of the core SAC protein kinetochore proximity association network. The Reactome circular interaction plot depicts the associations between the identified proteins within the core SAC protein proximity association network and the corresponding pathways in which they function. Legend presents color-coded pathways that correspond to the circular interaction plot.

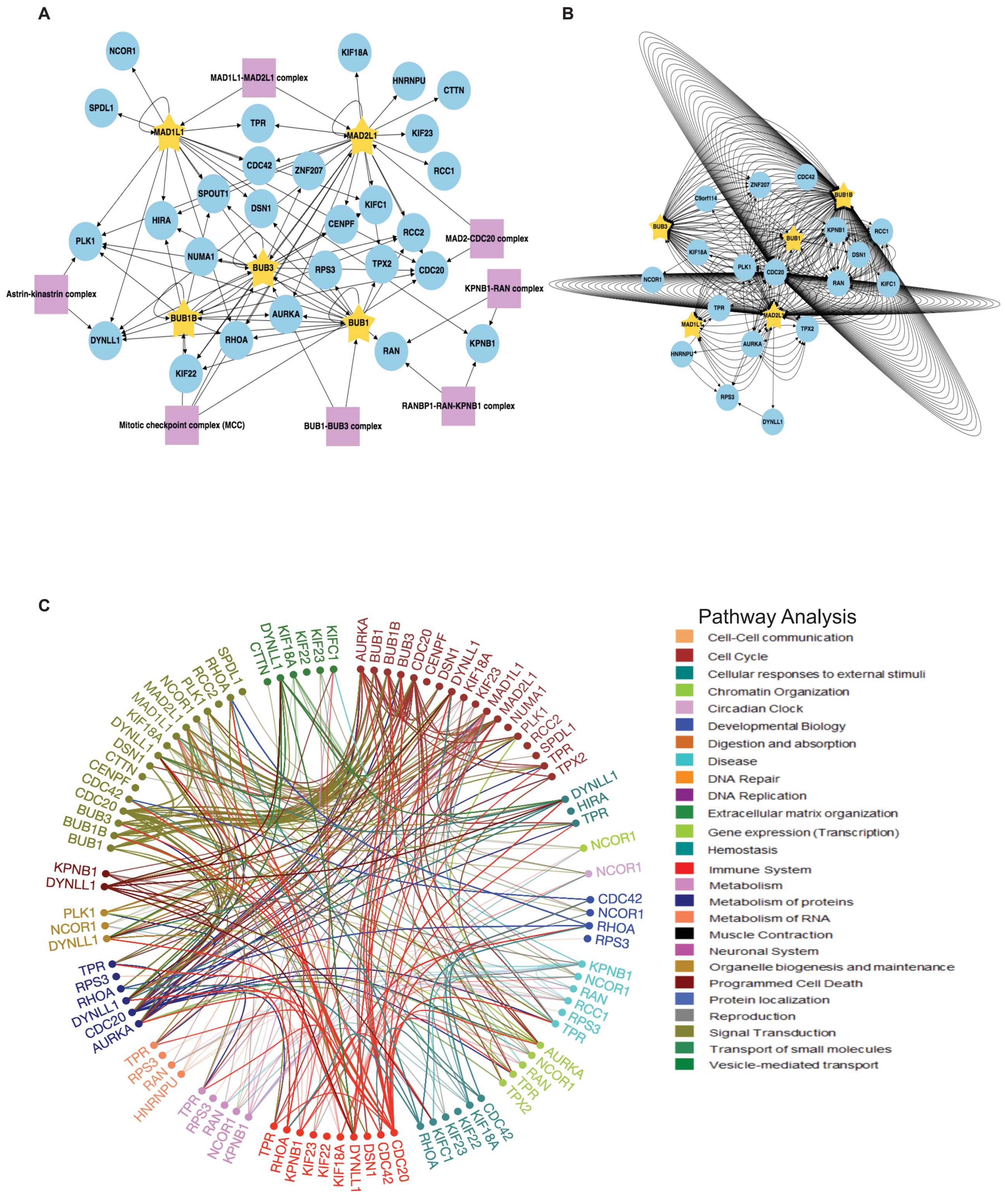

Supplemental Fig. S10. **The core SAC protein proximity association network using mitotic spindle related Gene Ontology annotations.** *A*, *B*, and *C* are as described in supplemental Fig. S9, except that the mitotic spindle related Gene Ontology annotations were applied to the analysis.

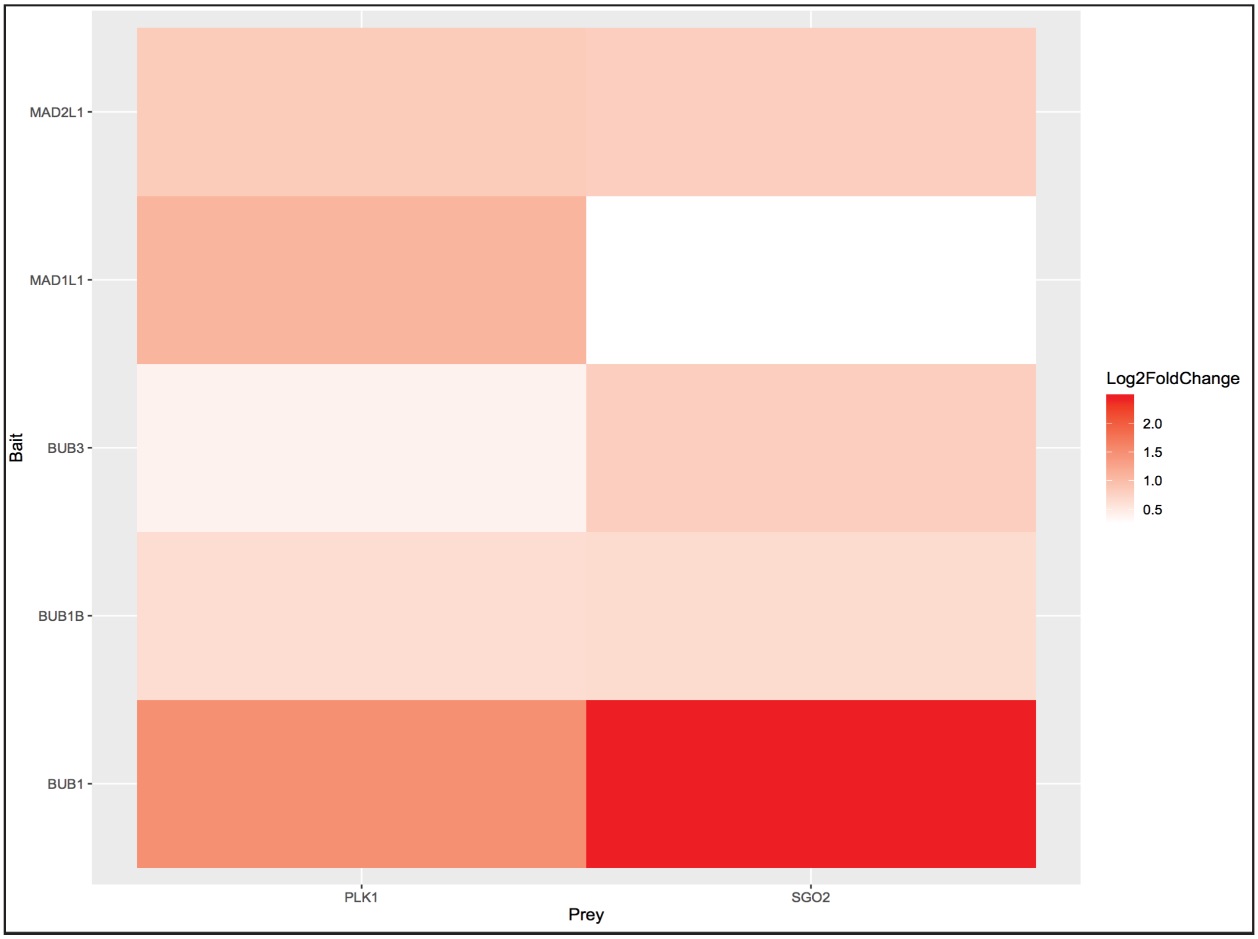

Supplemental Fig. S11. **BUB1 shows the strongest association with Astrin-Kinastrin minor complex components compared to other core SAC proteins.** The emPAI scores of the Astrin-Kinastrin minor complex components PLK1 and SGO2 in experimental purifications were compared against the control purification. The fold change was calculated and log2 transformed to visualize the relative change in test purifications. Note that both PLK1 and SGO2 showed a significant increase in the BUB1 purification. Also, note that the Astrin-Kinastrin minor complex component DYNLL1 was identified in all core SAC protein proximity association maps, but was excluded from this fold-change analysis as it was not identified the control purification.

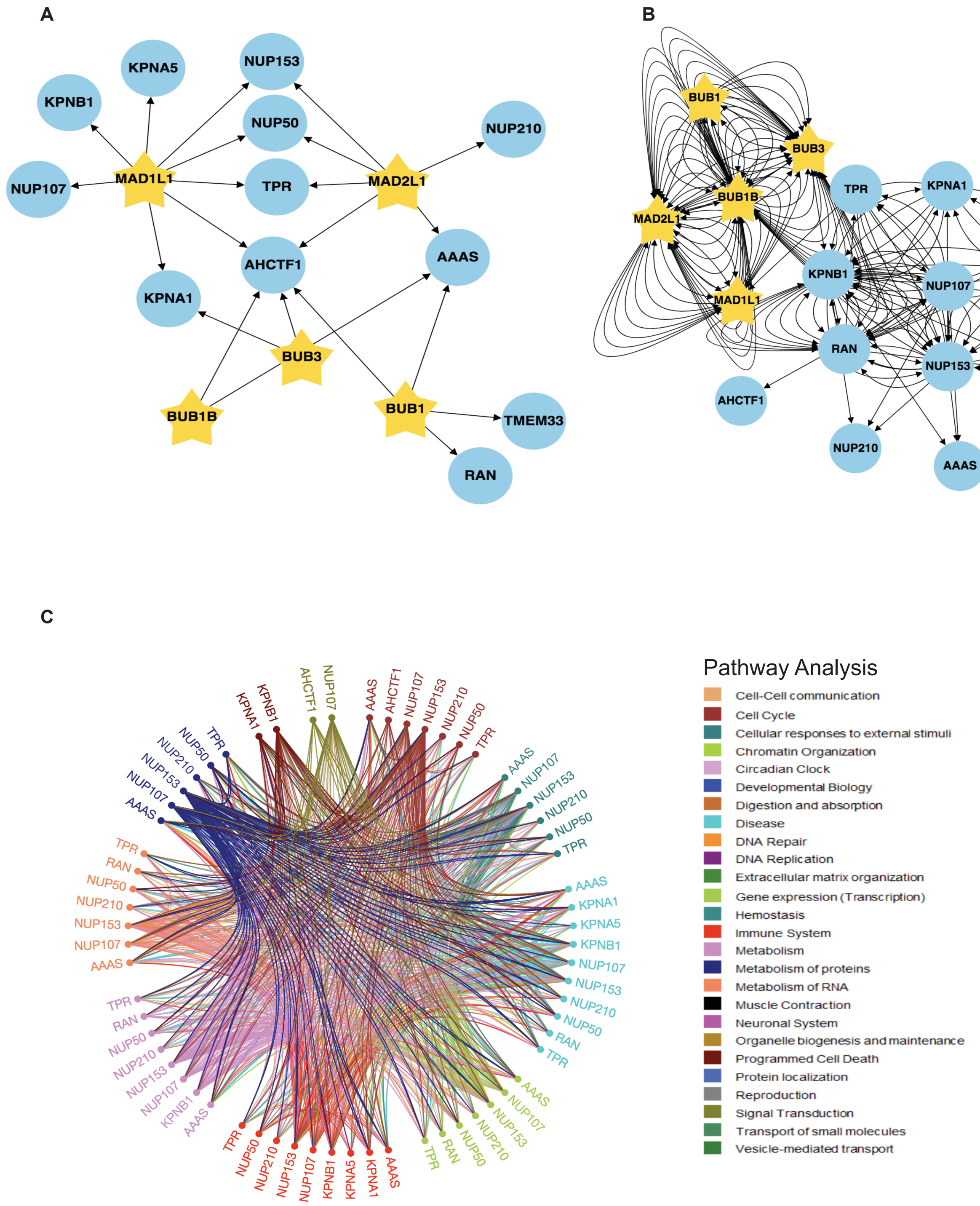

Supplemental Fig. S12. **The core SAC protein proximity association network using nuclear pore related Gene Ontology annotations.** *A*, *B*, and *C* are as described in supplemental Fig. S9, except that nuclear pore related Gene Ontology annotations were applied to the analysis.

**Supplemental Tables**

Supplemental Table S1. **List of reagents used.** List of reagents used and their pertinent information.

Supplemental Table S2. **List of primers used.** List of primers used to generate pGBioID2-27 and pGBioID2-47 vectors and pDONR221- *hBUB1, hBUB3, hBUBR1, hMAD1L1* and *hMAD2L1* vectors.

Supplemental Table S3. **List of vectors generated.** List of vectors that were generated and information on their properties.

Supplemental Table S4. **Summary of SAC protein proximity associated proteins.** Summary of BioID2-tagged BUB1, BUB3, BUBR1, MAD1L1 and MAD2L1 associated proteins identified by proximity labeling mass spectrometry analyses from mitotic cells. List includes protein UniProt accession number, emPAI score, and a description.

Supplemental Table S5. **List of Gene Ontology (GO) annotations used in the core SAC protein proximity association network analyses.** List of GO terms used in the Kinetochore, Mitotic Spindle, and Nuclear Pore network analyses.

Table S1

| Reagent | Company | Catalog Number | Description |
|---|---|---|---|
| Dynabeads™ MyOne™ Streptavidin C1 | Invitrogen | 65001 | Used as an affinity matrix during the purification of biotinylated proteins |
| TLA-100.3 tubes | Beckman | 349622 | Tubes for centrifuging protein lysates during the clearing step |
| QIAquick DNA gel extraction kit | Qiagen | 28704/28706 | Used in purifying PCR products from an agarose gel |
| BP clonase II | Invitrogen | 11789020 | Used for cloning ORF PCR products into the pDONR221 shuttle vector |
| LR clonase II | Invitrogen | 11791020 | Used for cloning the ORFs of genes of interest into the pGBioID2-tagging vector |
| *ccd* B Survival™ 2 T1R *E. coli* | Invitrogen | A10460 | Used for propgating shuttle vectors and pGBioID2 empty vectors |
| Fugene 6 | Promega | E2691 | Transfection reagent for transfecting vectors into human cells |
| Doxycycline | Clontech | 631311 | Drug for inducing Dox inducible protein expression |
| Hygromycin B | Invitrogen | 10687010 | Drug for selecting stable BioID2-tagged integrants |
| Kanamycin | Corning | 61-176-RG | Drug for selecting Kanamycin resistant bacterial colonies |
| Ampicillin | Fisher | BP1760-5 | Drug for selecting Ampicillin resistant bacterial colonies |
| 4-20% Tris Glycine SDS-PAGE gels | Biorad | 4561094 | Used for separating protein samples |
| Shuttle vector pDONR221 | Invitrogen | 12536017 | Shuttle vector for cloning the ORFs of genes of interest |
| Flippase expressing vector pOG44 | Invitrogen | V600520 | Vector that expresses the Flippase recombinase for integrating BioID2-tagged genes into the genome |
| Platinum Taq DNA Polymerase | ThermoFisher | 10966018 | Used for PCR amplification of the ORFs of genes of interest |
| 4X Laemmli sample buffer | Biorad | 1610747 | Protein sample buffer for SDS-PAGE |
| Lysogeny broth (LB) media | Fisher | BP9723-2 | Used for growing DH5α bacteria |
| DNA miniprep kit | Promega | A1222 | Used for making DNA plasmid minipreps |
| DMEM/F12 media | Hyclone | SH30023.01 | Used for growing HeLa human cells |
| FBS lacking Tet | Altanta Biologicals | S10350 | Used for making -Tet DMEM/F12 media for generating and growing inducible BioID2-tagged stable cell line |
| Trypsin | Hyclone | SH30042.01 | For lifting HeLa cell foci from plates |
| Protease inhibitor tablets | Roche | 11836170001 | Used for making protocol buffers, EDTA-free |
| 10% nonyl phenoxypolyethoxylethanol | Roche | 11332473001 | Used for making protocol buffers |
| PBS | Corning | 21-040-CM | Used for making protocol buffers |
| Triton-X-100 | Fisher | BP151-500 | Used for making protocol buffers |
| EDTA | ThermoFisher | 17892 | Used for making protocol buffers |
| EGTA | Acros Organics | 409911000 | Used for making protocol buffers |
| NaCl | Fisher | P217-3 | Used for making protocol buffers |
| KCl | Fisher | BP358-10 | Used for making protocol buffers |
| Dithiothreitol (DTT) | Fisher | BP172-25 | Used for making protocol buffers |
| MgCl2 | Fisher | M33-500 | Used for making protocol buffers |
| Tris base | Fisher | BP152-5 | Used for making protocol buffers |
| SDS | Fisher | BP166-500 | Used for making protocol buffers |
| Deoxycholic Acid | Sigma | D-6750 | Used for making protocol buffers |
| HEPES | Fisher | BP310-1 | Used for making protocol buffers |
| Lithium Chloride | Fisher | L120-500 | Used for making protocol buffers |
| 150 mm Tissue Culture Dish | ThermoFisher | 130183 | Used for maintaining and growing human cells |
| Biotin | Sigma | B4501-1G | Used for labeling cells in culture |
| Paclitaxel (Taxol) | Sigma | T7191-5MG | Used to arrest the cells in mitosis |
| BstBI | New England BioLabs | R0519S | Used for generating pGBioID2 vectors |
| AflII | New England BioLabs | R0520S | Used for generating pGBioID2 vectors |
| NheI | New England BioLabs | R0131S | Used for generating pGBioID2 vectors |
| XhoI | New England BioLabs | R0146S | Used for generating pGBioID2 vectors |
| BAY 1816032 inhibitor | MedChemExpress | HY-103020 | Used for inhibition of BUB1 kinase activity |
| PLK1 antibody | Abcam | ab17057 | Mouse anti-PLK1 antibody IF 1:100 |
| SGO2 anitbody | Bethyl | A301-262A-M | Rabbit anti-SGO2 antibody IF 1:100 |
| BioID2 antibody | BioFront Technologies | BID2-CP-100 | Chicken anti-BioID antibody WB 1:10000, IF 1:500 |
| GAPDH antibody | ProteinTech | 60004-1-1G | Mouse anti-GAPDH antibody WB 1:2000 |
| alpha-Tubulin antibody | Serotec | MCA77G | Rat anti-alpha-Tubulin antibody IF 1:500 |
| ACA antibody | Cortex Biochem | CS1058 | Human anti-ACA antibody IF 1:200 |
| BUB1 siRNA | Dharmacon | L-004102-00 | Used for BUB1 knock down experiments |
| Control siRNA | Dharmacon | D-001810-10 | Used as a control in knock down experiments |
| Lipofectamine RNAiMAX | Invitrogen | 13778-150 | siRNA transfection reagent |
| Opti-MEM I (1X) | Gibco | 31985-070 | siRNA transfection medium |

Table S2

| **Gene Name** | **Seq Name** |
|---|---|
| *BIRA-MYC-27* | BioID2. For |
| | BioID2. Rev |
| *BIRA-MYC-47* | BioID2. For |
| | BioID2. Rev |
| *BUB1* | pDONR223 |
| *BUB3* | pDONR221 |
| *BUBR1* | pDONR223 |
| *MAD1L1* | pDONR221 |
| *MAD2L1* | pDONR221 |

| **Annealing Sequence** |
| --- |
| ATGGAACAAAAACTCATCTCAGAAGAGGATCTCGACTTCAAGAACC |
| GCTTCTTCTCAGGCTGAACTCGCCGCT |
| ATGGAACAAAAACTCATCTCAGAAGAGGATCTCGACTTCAAGAACC |
| GCTTCTTCTCAGGCTGAACTCGCCGCT |

| Seq 5’ to 3’ |
|---|
| CGTTTAAACTTAAGATGGAACAAAAACTCATCTCAGAAGAGGATCTCGACTTCAAGAACC |
| GCTGGCGTTCGAATGAACCGCCACCACCTGAACCGCCACCACCTGAACCGCCACCACCTGAACC[illegible] |
| CGTTTAAACTTAAGATGGAACAAAAACTCATCTCAGAAGAGGATCTCGACTTCAAGAACC |
| GCTGGCGTTCGAAGCTTCTTCTCAGGCTGAACTCGCCGCT |
| DNASU: HsCD00399933 |
| DNASU: HsCD00043549 |
| DNASU: HsCD00399293 |
| Gift From Dr. Matthew Summers |
| DNASU: HsCD00296028 |

Table S3

| **Vector ID** | **Structure** | **Parental** | **Promote r** | **Bac Res** |
|---|---|---|---|---|
| pGBioID2-27 | BioID2-Myc-Linker 27 | pGLAP1 | CMV | Amp |
| pGBioID2-47 | BioID2-Myc-Linker 47 | pGLAP1 | CMV | Amp |
| pGBioID2-BUB1 | BioID2-Myc-Linker 47 | pGBioID2-47 | CMV | Amp |
| pGBioID2-BUB3 | BioID2-Myc-Linker 47 | pGBioID2-47 | CMV | Amp |
| pGBioID2-BUBR1 | BioID2-Myc-Linker 47 | pGBioID2-47 | CMV | Amp |
| pGBioID2-MAD1L1 | BioID2-Myc-Linker 47 | pGBioID2-47 | CMV | Amp |
| pGBioID2-MAD2L1 | BioID2-Myc-Linker 47 | pGBioID2-47 | CMV | Amp |

| Mam Res | Tet Regulated | Forward Sequencing Primer |
|---|---|---|
| Hyg | Yes | 5’-GGTGGGCCTGAGCGAGAAGGG-3’ |
| Hyg | Yes | 5’-GGTGGGCCTGAGCGAGAAGGG-3’ |
| Hyg | Yes | 5’-GGTGGGCCTGAGCGAGAAGGG-3’ |
| Hyg | Yes | 5’-GGTGGGCCTGAGCGAGAAGGG-3’ |
| Hyg | Yes | 5’-GGTGGGCCTGAGCGAGAAGGG-3’ |
| Hyg | Yes | 5’-GGTGGGCCTGAGCGAGAAGGG-3’ |
| Hyg | Yes | 5’-GGTGGGCCTGAGCGAGAAGGG-3’ |

| Reverse Sequencing Primer |
|---|
| 5’-AGCGGCGAGTTCAGCCTGAGAAGAAGC-3’ |
| 5’-AGCGGCGAGTTCAGCCTGAGAAGAAGC-3’ |
| 5’-AGCGGCGAGTTCAGCCTGAGAAGAAGC-3’ |
| 5’-AGCGGCGAGTTCAGCCTGAGAAGAAGC-3’ |
| 5’-AGCGGCGAGTTCAGCCTGAGAAGAAGC-3’ |
| 5’-AGCGGCGAGTTCAGCCTGAGAAGAAGC-3’ |
| 5’-AGCGGCGAGTTCAGCCTGAGAAGAAGC-3’ |

Table S4

| Accesion | emPAI | Description | Bait |
|---|---|---|---|
| A0A0A6YYL6 | 183 | Protein RPL1 | BUB1 |
| O00567 | 158.5 | Nucleolar pr | BUB1 |
| O00571 | 38.5 | ATP-depend | BUB1 |
| O00622 | 47 | Protein CYR6 | BUB1 |
| O43143 | 20.5 | Pre-mRNA-s | BUB1 |
| O43290 | 125.5 | U4/U6.U5 tr | BUB1 |
| O75151 | 22 | Lysine-specif | BUB1 |
| O75400 | 32.5 | Pre-mRNA-p | BUB1 |
| O75683 | 26 | Surfeit locus | BUB1 |
| O95299 | 22 | NADH dehyd | BUB1 |
| P04264 | 193 | Keratin, type | BUB1 |
| P04843 | 15.5 | Dolichyl-diph | BUB1 |
| P05141 | 85 | ADP/ATP tra | BUB1 |
| P07437 | 136 | Tubulin beta | BUB1 |
| P08195 | 52 | 4F2 cell-surf | BUB1 |
| P08670 | 142.5 | Vimentin OS | BUB1 |
| P09382 | 170.5 | Galectin-1 O | BUB1 |
| P10412 | 174.5 | Histone H1.4 | BUB1 |
| P10809 | 155.5 | 60 kDa heat | BUB1 |
| P11182 | 35 | Lipoamide a | BUB1 |
| P11387 | 165.5 | DNA topoiso | BUB1 |
| P11498 | 189.333333 | Pyruvate car | BUB1 |
| P12236 | 64.5 | ADP/ATP tra | BUB1 |
| P13995 | 13.5 | Bifunctional | BUB1 |
| P16401 | 187.5 | Histone H1.5 | BUB1 |
| P16615 | 18 | Sarcoplasmi | BUB1 |
| P17844 | 72.5 | Probable AT | BUB1 |
| P18583 | 19.5 | Protein SON | BUB1 |
| P22087 | 126 | rRNA 2'-O-m | BUB1 |
| P23284 | 84.5 | Peptidyl-pro | BUB1 |
| P23396 | 178.5 | 40S ribosom | BUB1 |
| P23528 | 38 | Cofilin-1 OS= | BUB1 |
| P25398 | 77 | 40S ribosom | BUB1 |
| P25705 | 111.666667 | ATP synthas | BUB1 |
| P27635 | 146.5 | 60S ribosom | BUB1 |
| P31943 | 130.5 | Heterogene | BUB1 |
| P35268 | 177.5 | 60S ribosom | BUB1 |
| P36542 | 51.5 | ATP synthas | BUB1 |
| P36578 | 176.5 | 60S ribosom | BUB1 |
| P37108 | 175.5 | Signal recog | BUB1 |
| P38646 | 32.5 | Stress-70 pr | BUB1 |

| | | | |
|---|---|---|---|
| P39023 | 167 | 60S ribosom | BUB1 |
| P46781 | 57.5 | 40S ribosom | BUB1 |
| P49411 | 127 | Elongation fa | BUB1 |
| P49458 | 158 | Signal recogi | BUB1 |
| P49756 | 53.5 | RNA-binding | BUB1 |
| P50454 | 93 | Serpin H1 OS | BUB1 |
| P51149 | 38.5 | Ras-related | BUB1 |
| P53350 | 42 | Serine/threo | BUB1 |
| P54198 | 31 | Protein HIRA | BUB1 |
| P57088 | 18.5 | Transmembr | BUB1 |
| P60709 | 186.5 | Actin, cytopl | BUB1 |
| P60866 | 157 | 40S ribosom | BUB1 |
| P61247 | 155 | 40S ribosom | BUB1 |
| P61254 | 180.5 | 60S ribosom | BUB1 |
| P61353 | 108.5 | 60S ribosom | BUB1 |
| P61513 | 155.5 | 60S ribosom | BUB1 |
| P61586 | 75.5 | Transforming | BUB1 |
| P62244 | 62.5 | 40S ribosom | BUB1 |
| P62249 | 100.5 | 40S ribosom | BUB1 |
| P62263 | 159 | 40S ribosom | BUB1 |
| P62280 | 96.5 | 40S ribosom | BUB1 |
| P62424 | 64 | 60S ribosom | BUB1 |
| P62701 | 164 | 40S ribosom | BUB1 |
| P62829 | 173.5 | 60S ribosom | BUB1 |
| P62847 | 182.5 | 40S ribosom | BUB1 |
| P62910 | 89.5 | 60S ribosom | BUB1 |
| P62913 | 118 | 60S ribosom | BUB1 |
| P62979 | 171 | Ubiquitin-40 | BUB1 |
| P63244 | 71 | Receptor of a | BUB1 |
| P82675 | 14 | 28S ribosom | BUB1 |
| P83731 | 139 | 60S ribosom | BUB1 |
| Q00325 | 17 | Phosphate ca | BUB1 |
| Q02878 | 65.5 | 60S ribosom | BUB1 |
| Q06830 | 89.5 | Peroxiredoxi | BUB1 |
| Q13085 | 199.333333 | Acetyl-CoA c | BUB1 |
| Q13162 | 36 | Peroxiredoxi | BUB1 |
| Q13523 | 98.5 | Serine/threo | BUB1 |
| Q13823 | 60.5 | Nucleolar GT | BUB1 |
| Q14331 | 59.5 | Protein FRG1 | BUB1 |
| Q14498 | 182 | RNA-binding | BUB1 |
| Q14684 | 179.5 | Ribosomal R | BUB1 |
| Q14807 | 63.5 | Kinesin-like | BUB1 |
| Q15149 | 67.5 | Plectin OS=H | BUB1 |

| | | | |
|---|---|---|---|
| Q15361 | 28 | Transcriptior | BUB1 |
| Q15365 | 34 | Poly(rC)-bind | BUB1 |
| Q15366 | 23 | Poly(rC)-bind | BUB1 |
| Q1ED39 | 43.5 | Lysine-rich n | BUB1 |
| Q2NL82 | 49 | Pre-rRNA-pr | BUB1 |
| Q3ZCQ8 | 32.5 | Mitochondria | BUB1 |
| Q562F6 | 169.5 | Shugoshin 2 | BUB1 |
| Q5FBB7 | 22.5 | Shugoshin 1 | BUB1 |
| Q5JTH9 | 20.5 | RRP12-like p | BUB1 |
| Q5QJE6 | 19 | Deoxynucleo | BUB1 |
| Q5T280 | 35 | Putative met | BUB1 |
| Q5T3I0 | 50 | G patch dom | BUB1 |
| Q69YH5 | 48 | Cell division | BUB1 |
| Q6DN12 | 21.5 | Multiple C2 a | BUB1 |
| Q6PCB5 | 49.5 | Round sperm | BUB1 |
| Q6UB99 | 8 | Ankyrin repe | BUB1 |
| Q6WKZ4 | 127.666667 | Rab11 family | BUB1 |
| Q7Z6E9 | 25 | E3 ubiquitin- | BUB1 |
| Q86WX3 | 57 | Active regula | BUB1 |
| Q86Y79 | 156 | Probable pep | BUB1 |
| Q8IY37 | 10.5 | Probable ATI | BUB1 |
| Q8IY81 | 71 | pre-rRNA pro | BUB1 |
| Q8NE71 | 69 | ATP-binding | BUB1 |
| Q8TBX8 | 35 | Phosphatidyl | BUB1 |
| Q8TCJ2 | 21 | Dolichyl-diph | BUB1 |
| Q8TDN6 | 26.5 | Ribosome bi | BUB1 |
| Q8WTT2 | 81.5 | Nucleolar co | BUB1 |
| Q8WYP5 | 32.5 | Protein ELYS | BUB1 |
| Q969Q0 | 43 | 60S ribosom | BUB1 |
| Q96GQ7 | 40 | Probable ATI | BUB1 |
| Q96IZ7 | 18 | Serine/Argin | BUB1 |
| Q99848 | 57.5 | Probable rRN | BUB1 |
| Q9BQ67 | 38.5 | Glutamate-ri | BUB1 |
| Q9BQG0 | 18 | Myb-binding | BUB1 |
| Q9BRT6 | 70.5 | Protein LLP h | BUB1 |
| Q9BSC4 | 27 | Nucleolar pro | BUB1 |
| Q9BU76 | 58.5 | Multiple mye | BUB1 |
| Q9BVJ6 | 29.5 | U3 small nuc | BUB1 |
| Q9BVP2 | 180 | Guanine nucl | BUB1 |
| Q9BXF6 | 38.5 | Rab11 family | BUB1 |
| Q9BXS0 | 19 | Collagen alpl | BUB1 |
| Q9BZE4 | 44 | Nucleolar GT | BUB1 |
| Q9BZF9 | 14.6666667 | Uveal autoar | BUB1 |

| | | | |
|---|---|---|---|
| Q9GZR7 | 49.5 | ATP-depende | BUB1 |
| Q9H0A0 | 34.3333333 | RNA cytidine | BUB1 |
| Q9H0S4 | 45 | Probable ATI | BUB1 |
| Q9H0U3 | 18 | Magnesium | BUB1 |
| Q9HCC0 | 78 | Methylcrotor | BUB1 |
| Q9HDC5 | 48 | Junctophilin- | BUB1 |
| Q9NP64 | 127.5 | Nucleolar pro | BUB1 |
| Q9NP72 | 22.5 | Ras-related | BUB1 |
| Q9NVP1 | 145 | ATP-depende | BUB1 |
| Q9NWB6 | 129 | Arginine and | BUB1 |
| Q9NZM5 | 53 | Ribosome bi | BUB1 |
| Q9P258 | 51 | Protein RCC2 | BUB1 |
| Q9UER7 | 22 | Death domai | BUB1 |
| Q9ULW0 | 47 | Targeting pro | BUB1 |
| Q9Y2X3 | 144.5 | Nucleolar pro | BUB1 |
| Q9Y305 | 26 | Acyl-coenzym | BUB1 |
| Q9Y3C1 | 103.5 | Nucleolar pro | BUB1 |
| Q9Y4P3 | 17.5 | Transducin b | BUB1 |
| A0A0A6YYL6 | 105.666667 | Protein RPL1 | BUB1B |
| O00567 | 84.3333333 | Nucleolar pro | BUB1B |
| O43290 | 55.5 | U4/U6.U5 tri | BUB1B |
| O60841 | 102.5 | Eukaryotic tr | BUB1B |
| O75151 | 21.5 | Lysine-specif | BUB1B |
| O75616 | 14.5 | GTPase Era, | BUB1B |
| O95232 | 68.5 | Luc7-like pro | BUB1B |
| O95299 | 26 | NADH dehyd | BUB1B |
| P05141 | 62.6666667 | ADP/ATP tra | BUB1B |
| P10809 | 76.6666667 | 60 kDa heat | BUB1B |
| P11387 | 130.5 | DNA topoiso | BUB1B |
| P13995 | 13.5 | Bifunctional | BUB1B |
| P15924 | 6 | Desmoplakin | BUB1B |
| P16403 | 149.5 | Histone H1.2 | BUB1B |
| P17844 | 52 | Probable ATI | BUB1B |
| P25398 | 35.5 | 40S ribosom | BUB1B |
| P25705 | 67.3333333 | ATP synthase | BUB1B |
| P27635 | 123 | 60S ribosom | BUB1B |
| P31943 | 76 | Heterogeneo | BUB1B |
| P35268 | 152.5 | 60S ribosom | BUB1B |
| P46013 | 42.5 | Proliferation | BUB1B |
| P46778 | 48.5 | 60S ribosom | BUB1B |
| P49411 | 114.333333 | Elongation fa | BUB1B |
| P49458 | 136.666667 | Signal recogi | BUB1B |
| P49756 | 37 | RNA-binding | BUB1B |

| | | | |
|---|---|---|---|
| P53350 | 23 | Serine/threo | BUB1B |
| P54198 | 19 | Protein HIRA | BUB1B |
| P61247 | 44.3333333 | 40S ribosom | BUB1B |
| P61254 | 149.333333 | 60S ribosom | BUB1B |
| P61586 | 50 | Transforming | BUB1B |
| P62241 | 157.666667 | 40S ribosom | BUB1B |
| P62263 | 163.666667 | 40S ribosom | BUB1B |
| P62266 | 132 | 40S ribosom | BUB1B |
| P62280 | 74 | 40S ribosom | BUB1B |
| P62701 | 69.3333333 | 40S ribosom | BUB1B |
| P62736 | 109 | Actin, aortic | BUB1B |
| P62829 | 100 | 60S ribosom | BUB1B |
| P62913 | 90 | 60S ribosom | BUB1B |
| P81605 | 63 | Dermcidin O | BUB1B |
| P82675 | 10.5 | 28S ribosom | BUB1B |
| Q00325 | 12.5 | Phosphate ca | BUB1B |
| Q02878 | 101.5 | 60S ribosom | BUB1B |
| Q03164 | 7 | Histone-lysin | BUB1B |
| Q05519 | 49 | Serine/argini | BUB1B |
| Q06830 | 71.6666667 | Peroxiredoxi | BUB1B |
| Q12873 | 10 | Chromodoma | BUB1B |
| Q13162 | 35.5 | Peroxiredoxi | BUB1B |
| Q13185 | 34 | Chromobox p | BUB1B |
| Q14684 | 139.5 | Ribosomal R | BUB1B |
| Q14807 | 58.5 | Kinesin-like p | BUB1B |
| Q15366 | 99 | Poly(rC)-bind | BUB1B |
| Q1ED39 | 16.5 | Lysine-rich n | BUB1B |
| Q3ZCQ8 | 14.3333333 | Mitochondria | BUB1B |
| Q562F6 | 104 | Shugoshin 2 | BUB1B |
| Q5JTH9 | 11.5 | RRP12-like p | BUB1B |
| Q5QJE6 | 23.5 | Deoxynucleo | BUB1B |
| Q5T280 | 22.3333333 | Putative met | BUB1B |
| Q5VTL8 | 23 | Pre-mRNA-s | BUB1B |
| Q66PJ3 | 41 | ADP-ribosyla | BUB1B |
| Q69YH5 | 53 | Cell division | BUB1B |
| Q6DN12 | 17.5 | Multiple C2 a | BUB1B |
| Q6PCB5 | 42.5 | Round sperm | BUB1B |
| Q6UB99 | 10.5 | Ankyrin repe | BUB1B |
| Q7Z6E9 | 18.5 | E3 ubiquitin- | BUB1B |
| Q86Y79 | 56 | Probable pep | BUB1B |
| Q8IWS0 | 44 | PHD finger p | BUB1B |
| Q8IY37 | 8 | Probable ATF | BUB1B |
| Q8NE71 | 63.5 | ATP-binding | BUB1B |

| | | | |
|---|---|---|---|
| Q8WYP5 | 43.5 | Protein ELYS | BUB1B |
| Q92922 | 8 | SWI/SNF cor | BUB1B |
| Q9BXF6 | 47.5 | Rab11 family | BUB1B |
| Q9BZE4 | 40 | Nucleolar GT | BUB1B |
| Q9BZF9 | 19 | Uveal autoar | BUB1B |
| Q9GZR7 | 36 | ATP-depende | BUB1B |
| Q9H0S4 | 13.5 | Probable ATI | BUB1B |
| Q9NVP1 | 69.5 | ATP-depende | BUB1B |
| Q9NWB6 | 76.6666667 | Arginine and | BUB1B |
| Q9P258 | 53.5 | Protein RCC2 | BUB1B |
| Q9P2D1 | 7.5 | Chromodoma | BUB1B |
| Q9UER7 | 27 | Death domai | BUB1B |
| Q9Y2X3 | 85 | Nucleolar pro | BUB1B |
| Q9Y305 | 28 | Acyl-coenzym | BUB1B |
| Q9Y4P3 | 10.5 | Transducin b | BUB1B |
| O43290 | 43 | U4/U6.U5 tri | BUB3 |
| P04843 | 15.5 | Dolichyl-diph | BUB3 |
| P10809 | 84.5 | 60 kDa heat | BUB3 |
| P16401 | 127 | Histone H1.5 | BUB3 |
| P16615 | 15.5 | Sarcoplasmic | BUB3 |
| P23528 | 49 | Cofilin-1 OS= | BUB3 |
| P31943 | 49 | Heterogeneo | BUB3 |
| P49458 | 126.5 | Signal recogi | BUB3 |
| P61254 | 146 | 60S ribosoma | BUB3 |
| P62266 | 131 | 40S ribosoma | BUB3 |
| P62829 | 63.5 | 60S ribosoma | BUB3 |
| P63244 | 48.5 | Receptor of a | BUB3 |
| P83731 | 115.5 | 60S ribosoma | BUB3 |
| Q06830 | 110.5 | Peroxiredoxi | BUB3 |
| Q13162 | 43 | Peroxiredoxi | BUB3 |
| Q13428 | 49 | Treacle prote | BUB3 |
| Q15365 | 28.5 | Poly(rC)-bind | BUB3 |
| Q15366 | 28 | Poly(rC)-bind | BUB3 |
| Q3ZCQ8 | 17.5 | Mitochondria | BUB3 |
| Q5T3I0 | 34.5 | G patch dom | BUB3 |
| Q6PCB5 | 29 | Round sperm | BUB3 |
| Q6WKZ4 | 138 | Rab11 family | BUB3 |
| Q7Z6E9 | 28.5 | E3 ubiquitin- | BUB3 |
| Q8WYP5 | 31.5 | Protein ELYS | BUB3 |
| Q969Q0 | 88 | 60S ribosoma | BUB3 |
| Q9BU76 | 58.5 | Multiple mye | BUB3 |
| Q9BZF9 | 17 | Uveal autoar | BUB3 |
| Q9NWB6 | 74 | Arginine and | BUB3 |

| | | | |
|---|---|---|---|
| Q9NX58 | 118.5 | Cell growth-r | BUB3 |
| A0A0A6YYL6 | 148.666667 | Protein RPL1 | MAD1L1 |
| E9PAV3 | 12.3333333 | Nascent poly | MAD1L1 |
| O00567 | 85 | Nucleolar pro | MAD1L1 |
| O14617 | 66.6666667 | AP-3 comple | MAD1L1 |
| O95232 | 92.5 | Luc7-like pro | MAD1L1 |
| O95299 | 26.5 | NADH dehyd | MAD1L1 |
| P05141 | 58.6666667 | ADP/ATP tra | MAD1L1 |
| P10809 | 102.666667 | 60 kDa heat s | MAD1L1 |
| P11182 | 20 | Lipoamide ac | MAD1L1 |
| P13995 | 13.5 | Bifunctional | MAD1L1 |
| P15924 | 7 | Desmoplakin | MAD1L1 |
| P17844 | 47.6666667 | Probable ATF | MAD1L1 |
| P23528 | 43.6666667 | Cofilin-1 OS= | MAD1L1 |
| P25398 | 54 | 40S ribosoma | MAD1L1 |
| P25705 | 80 | ATP synthase | MAD1L1 |
| P31327 | 131.333333 | Carbamoyl-p | MAD1L1 |
| P31943 | 49.6666667 | Heterogeneo | MAD1L1 |
| P36873 | 34 | Serine/threo | MAD1L1 |
| P39023 | 108 | 60S ribosoma | MAD1L1 |
| P49411 | 119.333333 | Elongation fa | MAD1L1 |
| P49458 | 94 | Signal recogn | MAD1L1 |
| P51149 | 46 | Ras-related p | MAD1L1 |
| P52292 | 15 | Importin sub | MAD1L1 |
| P54198 | 17.5 | Protein HIRA | MAD1L1 |
| P61247 | 48 | 40S ribosoma | MAD1L1 |
| P61254 | 156.333333 | 60S ribosoma | MAD1L1 |
| P61586 | 49 | Transforming | MAD1L1 |
| P62263 | 145.333333 | 40S ribosoma | MAD1L1 |
| P62280 | 57.6666667 | 40S ribosoma | MAD1L1 |
| P62701 | 106 | 40S ribosoma | MAD1L1 |
| P62829 | 68.6666667 | 60S ribosoma | MAD1L1 |
| P62913 | 73.6666667 | 60S ribosoma | MAD1L1 |
| P63244 | 42 | Receptor of a | MAD1L1 |
| P83731 | 105 | 60S ribosoma | MAD1L1 |
| Q00325 | 17.3333333 | Phosphate ca | MAD1L1 |
| Q06830 | 113 | Peroxiredoxin | MAD1L1 |
| Q13428 | 93.6666667 | Treacle prote | MAD1L1 |
| Q13523 | 109 | Serine/threo | MAD1L1 |
| Q13823 | 43 | Nucleolar GT | MAD1L1 |
| Q15365 | 28 | Poly(rC)-bind | MAD1L1 |
| Q1ED39 | 43 | Lysine-rich n | MAD1L1 |
| Q3ZCQ8 | 26.6666667 | Mitochondria | MAD1L1 |

| | | | |
|---|---|---|---|
| Q562F6 | 104.666667 | Shugoshin 2 | MAD1L1 |
| Q5T3I0 | 29 | G patch dom | MAD1L1 |
| Q66PJ3 | 28.5 | ADP-ribosyla | MAD1L1 |
| Q69YH5 | 64.3333333 | Cell division | MAD1L1 |
| Q6DN12 | 10 | Multiple C2 a | MAD1L1 |
| Q6PCB5 | 33.3333333 | Round sperm | MAD1L1 |
| Q6UB99 | 8.5 | Ankyrin repe | MAD1L1 |
| Q6WKZ4 | 122.666667 | Rab11 family | MAD1L1 |
| Q7Z6E9 | 16 | E3 ubiquitin- | MAD1L1 |
| Q8WYP5 | 57.6666667 | Protein ELYS | MAD1L1 |
| Q969Q0 | 75.5 | 60S ribosom | MAD1L1 |
| Q96IZ7 | 23 | Serine/Argin | MAD1L1 |
| Q9BRT6 | 57 | Protein LLP h | MAD1L1 |
| Q9BU76 | 40.6666667 | Multiple mye | MAD1L1 |
| Q9H0S4 | 17 | Probable ATI | MAD1L1 |
| Q9NP64 | 67.5 | Nucleolar pr | MAD1L1 |
| Q9NWB6 | 84.3333333 | Arginine and | MAD1L1 |
| Q9NX24 | 34.3333333 | H/ACA ribon | MAD1L1 |
| Q9NX58 | 127.666667 | Cell growth- | MAD1L1 |
| Q9Y2X3 | 92.3333333 | Nucleolar pr | MAD1L1 |
| Q9Y3C1 | 114 | Nucleolar pr | MAD1L1 |
| A0A0A6YYL6 | 150 | Protein RPL1 | MAD2L1 |
| O00567 | 116.333333 | Nucleolar pr | MAD2L1 |
| O00571 | 41.5 | ATP-depende | MAD2L1 |
| O00622 | 11.5 | Protein CYR6 | MAD2L1 |
| O14617 | 60 | AP-3 comple | MAD2L1 |
| O14641 | 18.3333333 | Segment pol | MAD2L1 |
| O15042 | 7 | U2 snRNP-as | MAD2L1 |
| O43143 | 28 | Pre-mRNA-s | MAD2L1 |
| O43290 | 80 | U4/U6.U5 tr | MAD2L1 |
| O60264 | 28.5 | SWI/SNF-rel | MAD2L1 |
| O75151 | 25.5 | Lysine-specif | MAD2L1 |
| O75400 | 40.5 | Pre-mRNA-p | MAD2L1 |
| O75616 | 18 | GTPase Era, | MAD2L1 |
| O76021 | 42 | Ribosomal L | MAD2L1 |
| O76094 | 60.5 | Signal recog | MAD2L1 |
| O95232 | 127 | Luc7-like pro | MAD2L1 |
| O95299 | 22 | NADH dehyd | MAD2L1 |
| O96028 | 17 | Histone-lysin | MAD2L1 |
| P04264 | 196.666667 | Keratin, type | MAD2L1 |
| P04843 | 15.5 | Dolichyl-diph | MAD2L1 |
| P05165 | 208.666667 | Propionyl-Co | MAD2L1 |
| P05166 | 161.5 | Propionyl-Co | MAD2L1 |

| | | | |
|---|---|---|---|
| P06748 | 35 | Nucleophosn | MAD2L1 |
| P07437 | 149 | Tubulin beta | MAD2L1 |
| P08195 | 52 | 4F2 cell-surf | MAD2L1 |
| P08238 | 16.5 | Heat shock p | MAD2L1 |
| P09382 | 133 | Galectin-1 O | MAD2L1 |
| P09874 | 96.5 | Poly [ADP-rib | MAD2L1 |
| P10412 | 195.666667 | Histone H1.4 | MAD2L1 |
| P10809 | 118 | 60 kDa heat s | MAD2L1 |
| P11021 | 28 | Endoplasmic | MAD2L1 |
| P11142 | 43 | Heat shock c | MAD2L1 |
| P11182 | 38.5 | Lipoamide a | MAD2L1 |
| P11387 | 161.5 | DNA topoiso | MAD2L1 |
| P11498 | 208 | Pyruvate carl | MAD2L1 |
| P12956 | 85 | X-ray repair c | MAD2L1 |
| P13010 | 150.5 | X-ray repair c | MAD2L1 |
| P16401 | 139.666667 | Histone H1.5 | MAD2L1 |
| P16403 | 203.333333 | Histone H1.2 | MAD2L1 |
| P16615 | 14.3333333 | Sarcoplasmic | MAD2L1 |
| P17480 | 105.333333 | Nucleolar tra | MAD2L1 |
| P17844 | 97.6666667 | Probable ATI | MAD2L1 |
| P19784 | 21.5 | Casein kinase | MAD2L1 |
| P22087 | 157.5 | rRNA 2'-O-m | MAD2L1 |
| P23284 | 81 | Peptidyl-prol | MAD2L1 |
| P23528 | 32 | Cofilin-1 OS= | MAD2L1 |
| P25398 | 62.5 | 40S ribosoma | MAD2L1 |
| P25705 | 118 | ATP synthase | MAD2L1 |
| P26368 | 9.5 | Splicing facto | MAD2L1 |
| P26373 | 207.666667 | 60S ribosoma | MAD2L1 |
| P27635 | 129.666667 | 60S ribosoma | MAD2L1 |
| P31327 | 120.666667 | Carbamoyl-p | MAD2L1 |
| P31943 | 60.6666667 | Heterogeneo | MAD2L1 |
| P35268 | 164 | 60S ribosoma | MAD2L1 |
| P35659 | 16.5 | Protein DEK ( | MAD2L1 |
| P36542 | 58 | ATP synthase | MAD2L1 |
| P36578 | 179.5 | 60S ribosoma | MAD2L1 |
| P36873 | 134 | Serine/threo | MAD2L1 |
| P37108 | 180.5 | Signal recogi | MAD2L1 |
| P38646 | 30.6666667 | Stress-70 pro | MAD2L1 |
| P39023 | 120 | 60S ribosoma | MAD2L1 |
| P46013 | 67.5 | Proliferation | MAD2L1 |
| P46781 | 49.5 | 40S ribosoma | MAD2L1 |
| P48426 | 15 | Phosphatidyl | MAD2L1 |
| P49411 | 122.333333 | Elongation fa | MAD2L1 |

| | | | |
|---|---|---|---|
| P49458 | 120 | Signal recogı | MAD2L1 |
| P49756 | 63 | RNA-binding | MAD2L1 |
| P51114 | 12.5 | Fragile X me | MAD2L1 |
| P51532 | 10 | Transcriptior | MAD2L1 |
| P52292 | 21.5 | Importin sub | MAD2L1 |
| P53350 | 27 | Serine/threo | MAD2L1 |
| P54198 | 28.5 | Protein HIRA | MAD2L1 |
| P60866 | 116.666667 | 40S ribosoma | MAD2L1 |
| P61247 | 67.3333333 | 40S ribosoma | MAD2L1 |
| P61254 | 175.666667 | 60S ribosoma | MAD2L1 |
| P61353 | 152 | 60S ribosoma | MAD2L1 |
| P61586 | 51 | Transforming | MAD2L1 |
| P62241 | 165.333333 | 40S ribosoma | MAD2L1 |
| P62244 | 63.5 | 40S ribosoma | MAD2L1 |
| P62249 | 91 | 40S ribosoma | MAD2L1 |
| P62263 | 188.666667 | 40S ribosoma | MAD2L1 |
| P62266 | 155.666667 | 40S ribosoma | MAD2L1 |
| P62280 | 66 | 40S ribosoma | MAD2L1 |
| P62424 | 50.5 | 60S ribosoma | MAD2L1 |
| P62701 | 123.666667 | 40S ribosoma | MAD2L1 |
| P62736 | 153 | Actin, aortic | MAD2L1 |
| P62750 | 151.666667 | 60S ribosoma | MAD2L1 |
| P62805 | 109 | Histone H4 C | MAD2L1 |
| P62829 | 103.666667 | 60S ribosoma | MAD2L1 |
| P62847 | 187 | 40S ribosoma | MAD2L1 |
| P62854 | 172.333333 | 40S ribosoma | MAD2L1 |
| P62899 | 65.5 | 60S ribosoma | MAD2L1 |
| P62910 | 71 | 60S ribosoma | MAD2L1 |
| P62913 | 126 | 60S ribosoma | MAD2L1 |
| P62917 | 183.5 | 60S ribosoma | MAD2L1 |
| P62979 | 166 | Ubiquitin-40S | MAD2L1 |
| P63244 | 42 | Receptor of a | MAD2L1 |
| P68363 | 98 | Tubulin alpha | MAD2L1 |
| P81605 | 47 | Dermcidin OS | MAD2L1 |
| P82675 | 34.5 | 28S ribosoma | MAD2L1 |
| P83731 | 136 | 60S ribosoma | MAD2L1 |
| Q00325 | 17 | Phosphate ca | MAD2L1 |
| Q02809 | 8 | Procollagen-l | MAD2L1 |
| Q02878 | 59 | 60S ribosoma | MAD2L1 |
| Q02880 | 37 | DNA topoisoı | MAD2L1 |
| Q03164 | 13 | Histone-lysin | MAD2L1 |
| Q06830 | 69 | Peroxiredoxiı | MAD2L1 |
| Q07065 | 136 | Cytoskeleton | MAD2L1 |

| | | | |
|---|---|---|---|
| Q12873 | 13.5 | Chromodoma | MAD2L1 |
| Q12931 | 15 | Heat shock p | MAD2L1 |
| Q13085 | 207.333333 | Acetyl-CoA c | MAD2L1 |
| Q13162 | 23 | Peroxiredoxi | MAD2L1 |
| Q13185 | 73 | Chromobox p | MAD2L1 |
| Q13427 | 45.5 | Peptidyl-prol | MAD2L1 |
| Q13428 | 57.3333333 | Treacle prote | MAD2L1 |
| Q13523 | 113 | Serine/threo | MAD2L1 |
| Q13823 | 55.5 | Nucleolar GT | MAD2L1 |
| Q14331 | 59 | Protein FRG1 | MAD2L1 |
| Q14498 | 126.666667 | RNA-binding | MAD2L1 |
| Q14684 | 176 | Ribosomal R | MAD2L1 |
| Q14807 | 66.5 | Kinesin-like p | MAD2L1 |
| Q15149 | 49 | Plectin OS=H | MAD2L1 |
| Q15366 | 28.3333333 | Poly(rC)-bind | MAD2L1 |
| Q1ED39 | 15.3333333 | Lysine-rich n | MAD2L1 |
| Q2NL82 | 50.5 | Pre-rRNA-pro | MAD2L1 |
| Q3ZCQ8 | 24.3333333 | Mitochondria | MAD2L1 |
| Q562F6 | 132.333333 | Shugoshin 2 | MAD2L1 |
| Q5JTH9 | 20.5 | RRP12-like p | MAD2L1 |
| Q5QJE6 | 19 | Deoxynucleo | MAD2L1 |
| Q5SSJ5 | 40.5 | Heterochrom | MAD2L1 |
| Q5T3I0 | 33 | G patch dom | MAD2L1 |
| Q5VTL8 | 32 | Pre-mRNA-s | MAD2L1 |
| Q66PJ3 | 28 | ADP-ribosyla | MAD2L1 |
| Q69YH5 | 49.6666667 | Cell division | MAD2L1 |
| Q6DN12 | 16.3333333 | Multiple C2 a | MAD2L1 |
| Q6PCB5 | 18.6666667 | Round sperm | MAD2L1 |
| Q6UB99 | 11 | Ankyrin repe | MAD2L1 |
| Q6WKZ4 | 129.333333 | Rab11 family | MAD2L1 |
| Q7L014 | 133 | Probable ATP | MAD2L1 |
| Q7Z6E9 | 19.5 | E3 ubiquitin- | MAD2L1 |
| Q7Z7K6 | 17 | Centromere | MAD2L1 |
| Q86UE4 | 77.5 | Protein LYRIC | MAD2L1 |
| Q86WX3 | 48 | Active regula | MAD2L1 |
| Q86Y79 | 116 | Probable pep | MAD2L1 |
| Q8IY37 | 10.5 | Probable ATP | MAD2L1 |
| Q8NE71 | 85 | ATP-binding | MAD2L1 |
| Q8NEV1 | 20 | Casein kinase | MAD2L1 |
| Q8TAA9 | 20.5 | Vang-like pro | MAD2L1 |
| Q8TCJ2 | 19 | Dolichyl-diph | MAD2L1 |
| Q8TDD1 | 68.5 | ATP-depende | MAD2L1 |
| Q8TDN6 | 22 | Ribosome bi | MAD2L1 |

| | | | |
|---|---|---|---|
| Q8WTT2 | 65 | Nucleolar co | MAD2L1 |
| Q8WX93 | 5.66666667 | Palladin OS= | MAD2L1 |
| Q8WXA9 | 31.5 | Splicing regu | MAD2L1 |
| Q8WY36 | 35.5 | HMG box tra | MAD2L1 |
| Q8WYP5 | 43.6666667 | Protein ELYS | MAD2L1 |
| Q92997 | 20.3333333 | Segment pol | MAD2L1 |
| Q96AG4 | 49 | Leucine-rich | MAD2L1 |
| Q96RQ3 | 192 | Methylcrotor | MAD2L1 |
| Q99848 | 49.5 | Probable rRN | MAD2L1 |
| Q9BQ67 | 59 | Glutamate-ri | MAD2L1 |
| Q9BQG0 | 11 | Myb-binding | MAD2L1 |
| Q9BRT6 | 74 | Protein LLP h | MAD2L1 |
| Q9BSC4 | 27 | Nucleolar pro | MAD2L1 |
| Q9BVJ6 | 29.5 | U3 small nuc | MAD2L1 |
| Q9BVP2 | 211 | Guanine nucl | MAD2L1 |
| Q9BXF6 | 51.5 | Rab11 family | MAD2L1 |
| Q9BZE4 | 37.6666667 | Nucleolar GT | MAD2L1 |
| Q9BZF9 | 15 | Uveal autoar | MAD2L1 |
| Q9GZR7 | 35.6666667 | ATP-depende | MAD2L1 |
| Q9H0A0 | 30.3333333 | RNA cytidine | MAD2L1 |
| Q9H0S4 | 33 | Probable ATP | MAD2L1 |
| Q9H8G2 | 18 | Caspase acti | MAD2L1 |
| Q9HCC0 | 144.5 | Methylcrotor | MAD2L1 |
| Q9NP72 | 22.5 | Ras-related p | MAD2L1 |
| Q9NRL2 | 12 | Bromodomai | MAD2L1 |
| Q9NVP1 | 61 | ATP-depende | MAD2L1 |
| Q9NWB6 | 112 | Arginine and | MAD2L1 |
| Q9NX58 | 111.666667 | Cell growth-r | MAD2L1 |
| Q9P0L0 | 55 | Vesicle-asso | MAD2L1 |
| Q9P258 | 54.5 | Protein RCC2 | MAD2L1 |
| Q9P2D1 | 14 | Chromodoma | MAD2L1 |
| Q9UER7 | 29.5 | Death domai | MAD2L1 |
| Q9UHB9 | 11.5 | Signal recogr | MAD2L1 |
| Q9UIG0 | 45 | Tyrosine-pro | MAD2L1 |
| Q9ULW0 | 40.6666667 | Targeting pro | MAD2L1 |
| Q9Y2X3 | 118.666667 | Nucleolar pro | MAD2L1 |
| Q9Y305 | 25.5 | Acyl-coenzym | MAD2L1 |
| Q9Y383 | 169.5 | Putative RNA | MAD2L1 |
| Q9Y3C1 | 120 | Nucleolar pro | MAD2L1 |
| Q9Y4P3 | 10.5 | Transducin b | MAD2L1 |
| Q9Y6C9 | 21 | Mitochondria | MAD2L1 |
| A0FGR8 | 16 | Extended syr | BUB1 |
| H7C0C1 | 19 | Uncharacteri | BUB1 |

| | | | |
|---|---|---|---|
| O14965 | 23.5 | Aurora kinas | BUB1 |
| O15427 | 7 | Monocarboxy | BUB1 |
| O43670 | 6 | BUB3-intera | BUB1 |
| O43683 | 90 | Mitotic checl | BUB1 |
| O43684 | 61.5 | Mitotic checl | BUB1 |
| O95433 | 18 | Activator of ! | BUB1 |
| O95639 | 11 | Cleavage and | BUB1 |
| O95782 | 4.5 | AP-2 comple | BUB1 |
| P06858 | 47 | Lipoprotein li | BUB1 |
| P07203 | 23.5 | Glutathione | BUB1 |
| P15880 | 11 | 40S ribosoma | BUB1 |
| P29375 | 6.5 | Lysine-specif | BUB1 |
| P35240 | 7.5 | Merlin OS=H | BUB1 |
| P35908 | 169.5 | Keratin, type | BUB1 |
| P42677 | 104 | 40S ribosoma | BUB1 |
| P46779 | 68.5 | 60S ribosoma | BUB1 |
| P46977 | 8 | Dolichyl-diph | BUB1 |
| P50402 | 53 | Emerin OS=H | BUB1 |
| P51571 | 125 | Translocon-a | BUB1 |
| P53985 | 6 | Monocarboxy | BUB1 |
| P60842 | 15 | Eukaryotic in | BUB1 |
| P61019 | 22 | Ras-related | BUB1 |
| P61106 | 54.5 | Ras-related | BUB1 |
| P63151 | 9.5 | Serine/threo | BUB1 |
| P63167 | 51 | Dynein light | BUB1 |
| Q07157 | 14.5 | Tight junctio | BUB1 |
| Q12797 | 12 | Aspartyl/asp | BUB1 |
| Q13454 | 8 | Tumor suppr | BUB1 |
| Q5JWF2 | 3 | Guanine nucl | BUB1 |
| Q6NZI2 | 25 | Caveolae-ass | BUB1 |
| Q6PJG2 | 6 | ELM2 and SA | BUB1 |
| Q6ZNB6 | 12.5 | NF-X1-type z | BUB1 |
| Q8NG31 | 12 | Kinetochore | BUB1 |
| Q8TA86 | 50.5 | Retinitis pigr | BUB1 |
| Q8WWC4 | 10 | m-AAA prote | BUB1 |
| Q96A33 | 19 | Coiled-coil do | BUB1 |
| Q96EY4 | 22 | Translation r | BUB1 |
| Q96HE9 | 27 | Proline-rich | BUB1 |
| Q96KM6 | 5 | Zinc finger p | BUB1 |
| Q9BQ48 | 34 | 39S ribosoma | BUB1 |
| Q9BQE3 | 60 | Tubulin alpha | BUB1 |
| Q9BZF3 | 8 | Oxysterol-bir | BUB1 |
| Q9H2Y7 | 2 | Zinc finger p | BUB1 |

| | | | |
|---|---|---|---|
| Q9H410 | 8 | Kinetochore- | BUB1 |
| Q9H4L5 | 28.5 | Oxysterol-bir | BUB1 |
| Q9H8M2 | 27 | Bromodomai | BUB1 |
| Q9NPF2 | 8 | Carbohydrate | BUB1 |
| Q9NPG3 | 11 | Ubinuclein-1 | BUB1 |
| Q9NSE4 | 3 | Isoleucine--t | BUB1 |
| Q9NX20 | 12 | 39S ribosom | BUB1 |
| Q9NY93 | 30.5 | Probable ATI | BUB1 |
| Q9NZ01 | 24.5 | Very-long-ch | BUB1 |
| Q9UGP8 | 8 | Translocatio | BUB1 |
| Q9UHI8 | 14.5 | A disintegrin | BUB1 |
| Q9UL25 | 14 | Ras-related | BUB1 |
| Q9UM01 | 6 | Y+L amino a | BUB1 |
| Q9Y3U8 | 28 | 60S ribosom | BUB1 |
| Q9Y4F1 | 4.5 | FERM, ARHG | BUB1 |
| A0FGR8 | 6 | Extended syr | BUB1B |
| O00763 | 28 | Acetyl-CoA c | BUB1B |
| O15427 | 7 | Monocarboxy | BUB1B |
| O43670 | 6 | BUB3-intera | BUB1B |
| O43684 | 21.6666667 | Mitotic checl | BUB1B |
| O60566 | 12.3333333 | Mitotic checl | BUB1B |
| O95292 | 33 | Vesicle-asso | BUB1B |
| P14923 | 14.5 | Junction plak | BUB1B |
| P15880 | 11 | 40S ribosom | BUB1B |
| P29375 | 6 | Lysine-specif | BUB1B |
| P31930 | 6 | Cytochrome | BUB1B |
| P35030 | 13.6666667 | Trypsin-3 OS | BUB1B |
| P35908 | 137.5 | Keratin, type | BUB1B |
| P47914 | 30 | 60S ribosom | BUB1B |
| P50402 | 24 | Emerin OS=H | BUB1B |
| P51151 | 15 | Ras-related | BUB1B |
| P51571 | 56 | Translocon-a | BUB1B |
| P53985 | 6 | Monocarboxy | BUB1B |
| P60842 | 7 | Eukaryotic in | BUB1B |
| P62140 | 33.5 | Serine/threo | BUB1B |
| P62820 | 80 | Ras-related | BUB1B |
| P63151 | 6 | Serine/threo | BUB1B |
| P68366 | 63 | Tubulin alph | BUB1B |
| P68371 | 87.5 | Tubulin beta | BUB1B |
| Q02413 | 9 | Desmoglein- | BUB1B |
| Q08554 | 3 | Desmocollin | BUB1B |
| Q12797 | 4 | Aspartyl/asp | BUB1B |
| Q12830 | 1.5 | Nucleosome | BUB1B |

| | | | |
|---|---|---|---|
| Q12834 | 6 | Cell division ( | BUB1B |
| Q14839 | 4.5 | Chromodoma | BUB1B |
| Q6PJG2 | 12 | ELM2 and SA | BUB1B |
| Q6ZNB6 | 11 | NF-X1-type z | BUB1B |
| Q70SY1 | 12 | Cyclic AMP-r | BUB1B |
| Q71UM5 | 93 | 40S ribosom | BUB1B |
| Q8WXI9 | 13 | Transcriptior | BUB1B |
| Q96EY4 | 14 | Translation r | BUB1B |
| Q96HE9 | 8 | Proline-rich p | BUB1B |
| Q96KM6 | 3 | Zinc finger p | BUB1B |
| Q96QE3 | 2 | ATPase fami | BUB1B |
| Q9BQ39 | 4 | ATP-depende | BUB1B |
| Q9H2Y7 | 4.5 | Zinc finger p | BUB1B |
| Q9H4L5 | 24.5 | Oxysterol-bir | BUB1B |
| Q9H8M2 | 10 | Bromodomai | BUB1B |
| Q9NPF2 | 8 | Carbohydrate | BUB1B |
| Q9NPG3 | 6.5 | Ubinuclein-1 | BUB1B |
| Q9NSE4 | 3 | Isoleucine--t | BUB1B |
| Q9NXE8 | 7 | Pre-mRNA-s | BUB1B |
| Q9NY93 | 19.5 | Probable ATI | BUB1B |
| Q9NYP7 | 9 | Elongation o | BUB1B |
| Q9P031 | 12 | Thyroid trans | BUB1B |
| Q9UGP8 | 6 | Translocatior | BUB1B |
| Q9UGU5 | 7.5 | HMG domair | BUB1B |
| Q9UJS0 | 6.5 | Calcium-binc | BUB1B |
| Q9UL25 | 14 | Ras-related p | BUB1B |
| Q9UNX3 | 160 | 60S ribosom | BUB1B |
| Q9UPQ3 | 3 | Arf-GAP with | BUB1B |
| O43670 | 24.5 | BUB3-intera | BUB3 |
| O43683 | 102.5 | Mitotic checl | BUB3 |
| O43684 | 126 | Mitotic checl | BUB3 |
| O60293 | 19.5 | Zinc finger C | BUB3 |
| O60566 | 24.5 | Mitotic checl | BUB3 |
| P27824 | 5 | Calnexin OS= | BUB3 |
| P47914 | 41.5 | 60S ribosom | BUB3 |
| P50402 | 17.5 | Emerin OS=H | BUB3 |
| P51571 | 37 | Translocon-a | BUB3 |
| Q12834 | 12.5 | Cell division ( | BUB3 |
| Q5VWQ0 | 9 | Lysine-specif | BUB3 |
| Q8NG31 | 23 | Kinetochore s | BUB3 |
| Q9BQE3 | 112.5 | Tubulin alpha | BUB3 |
| Q9P2N5 | 42 | RNA-binding | BUB3 |
| Q9Y4F1 | 3 | FERM, ARHG | BUB3 |

| | | | |
|---|---|---|---|
| A0A075B6Z2 | 154 | T cell recept | MAD1L1 |
| B4DLN1 | 7 | cDNA FLJ601 | MAD1L1 |
| H3BNC9 | 5 | Uncharacteri | MAD1L1 |
| O15131 | 8 | Importin sub | MAD1L1 |
| P00966 | 7 | Argininosucc | MAD1L1 |
| P06733 | 14.5 | Alpha-enolas | MAD1L1 |
| P06858 | 6 | Lipoprotein l | MAD1L1 |
| P12270 | 55.3333333 | Nucleoprotei | MAD1L1 |
| P14923 | 10 | Junction plak | MAD1L1 |
| P15880 | 14.6666667 | 40S ribosoma | MAD1L1 |
| P20290 | 23.5 | Transcriptior | MAD1L1 |
| P34931 | 5 | Heat shock 7 | MAD1L1 |
| P35908 | 149 | Keratin, type | MAD1L1 |
| P36969 | 23.5 | Phospholipid | MAD1L1 |
| P46779 | 22 | 60S ribosoma | MAD1L1 |
| P47914 | 72.3333333 | 60S ribosoma | MAD1L1 |
| P49790 | 57.6666667 | Nuclear pore | MAD1L1 |
| P51151 | 15 | Ras-related | MAD1L1 |
| P51571 | 31.3333333 | Translocon-a | MAD1L1 |
| P57740 | 7.33333333 | Nuclear pore | MAD1L1 |
| P60842 | 15 | Eukaryotic in | MAD1L1 |
| P63151 | 9.5 | Serine/threo | MAD1L1 |
| P63167 | 65 | Dynein light | MAD1L1 |
| P68366 | 100.666667 | Tubulin alpha | MAD1L1 |
| P68371 | 118.333333 | Tubulin beta | MAD1L1 |
| Q07157 | 7 | Tight junctio | MAD1L1 |
| Q14839 | 3 | Chromodoma | MAD1L1 |
| Q15084 | 7 | Protein disul | MAD1L1 |
| Q15147 | 2 | 1-phosphatic | MAD1L1 |
| Q2VIR3 | 6 | Eukaryotic tr | MAD1L1 |
| Q5VWQ0 | 5.5 | Lysine-specif | MAD1L1 |
| Q5W0B1 | 17 | RING finger | MAD1L1 |
| Q6NZI2 | 8 | Caveolae-ass | MAD1L1 |
| Q6ZNB6 | 6 | NF-X1-type z | MAD1L1 |
| Q71UM5 | 52.5 | 40S ribosoma | MAD1L1 |
| Q86UY6 | 12 | N-alpha-acet | MAD1L1 |
| Q8NG31 | 8 | Kinetochore | MAD1L1 |
| Q8TA86 | 49.5 | Retinitis pigr | MAD1L1 |
| Q96A33 | 6 | Coiled-coil d | MAD1L1 |
| Q96EA4 | 5 | Protein Spinc | MAD1L1 |
| Q96EY4 | 22 | Translation r | MAD1L1 |
| Q96HE9 | 8 | Proline-rich | MAD1L1 |
| Q9BZF3 | 4.5 | Oxysterol-bir | MAD1L1 |

| | | | |
|---|---|---|---|
| Q9H2Y7 | 2 | Zinc finger p | MAD1L1 |
| Q9H4L5 | 10 | Oxysterol-bir | MAD1L1 |
| Q9H8M2 | 15.5 | Bromodoma | MAD1L1 |
| Q9NPG3 | 4 | Ubinuclein-1 | MAD1L1 |
| Q9P0U3 | 12.3333333 | Sentrin-speci | MAD1L1 |
| Q9P2N5 | 15.6666667 | RNA-binding | MAD1L1 |
| Q9UHF7 | 7 | Zinc finger tr | MAD1L1 |
| Q9UJS0 | 4 | Calcium-binc | MAD1L1 |
| Q9UKX7 | 94 | Nuclear pore | MAD1L1 |
| Q9UNX9 | 7 | ATP-sensitiv | MAD1L1 |
| Q9UQP3 | 2 | Tenascin-N ( | MAD1L1 |
| Q9Y6D9 | 162.333333 | Mitotic spinc | MAD1L1 |
| A0FGR8 | 8 | Extended syr | MAD2L1 |
| A0JLT2 | 27.5 | Mediator of | MAD2L1 |
| H7C0C1 | 12 | Uncharacteri | MAD2L1 |
| O00411 | 6 | DNA-directe | MAD2L1 |
| O00541 | 5 | Pescadillo hc | MAD2L1 |
| O14647 | 2 | Chromodoma | MAD2L1 |
| O14965 | 11 | Aurora kinas | MAD2L1 |
| O15427 | 7 | Monocarbox | MAD2L1 |
| O43670 | 6 | BUB3-intera | MAD2L1 |
| O43684 | 23.5 | Mitotic checl | MAD2L1 |
| O60831 | 17 | PRA1 family | MAD2L1 |
| O95235 | 3 | Kinesin-like p | MAD2L1 |
| O95292 | 33 | Vesicle-asso | MAD2L1 |
| O95478 | 24 | Ribosome bi | MAD2L1 |
| O95782 | 16.5 | AP-2 comple | MAD2L1 |
| P00966 | 7 | Argininosucc | MAD2L1 |
| P04908 | 164.5 | Histone H2A | MAD2L1 |
| P06858 | 13 | Lipoprotein l | MAD2L1 |
| P09038 | 10.5 | Fibroblast gr | MAD2L1 |
| P12270 | 12.3333333 | Nucleoprotei | MAD2L1 |
| P14618 | 6 | Pyruvate kina | MAD2L1 |
| P15880 | 11 | 40S ribosom | MAD2L1 |
| P18077 | 27 | 60S ribosom | MAD2L1 |
| P21127 | 9.5 | Cyclin-depen | MAD2L1 |
| P29375 | 7.5 | Lysine-specif | MAD2L1 |
| P30048 | 12 | Thioredoxin- | MAD2L1 |
| P31930 | 6 | Cytochrome | MAD2L1 |
| P35030 | 10 | Trypsin-3 OS | MAD2L1 |
| P35908 | 191 | Keratin, type | MAD2L1 |
| P46779 | 34.5 | 60S ribosom | MAD2L1 |
| P49790 | 4 | Nuclear pore | MAD2L1 |

| | | | |
|---|---|---|---|
| P50402 | 31 | Emerin OS=H | MAD2L1 |
| P51571 | 120.5 | Translocon-a | MAD2L1 |
| P60842 | 19 | Eukaryotic in | MAD2L1 |
| P62081 | 42 | 40S ribosoma | MAD2L1 |
| P62140 | 138.5 | Serine/threo | MAD2L1 |
| P63010 | 13.5 | AP-2 comple | MAD2L1 |
| P63151 | 9.5 | Serine/threo | MAD2L1 |
| P68371 | 181 | Tubulin beta- | MAD2L1 |
| P78527 | 4 | DNA-depend | MAD2L1 |
| Q07157 | 14.6666667 | Tight junctio | MAD2L1 |
| Q12797 | 16.5 | Aspartyl/asp | MAD2L1 |
| Q12830 | 2 | Nucleosome- | MAD2L1 |
| Q13257 | 108.333333 | Mitotic spinc | MAD2L1 |
| Q13501 | 23 | Sequestosom | MAD2L1 |
| Q14562 | 2 | ATP-depende | MAD2L1 |
| Q14739 | 5 | Lamin-B rece | MAD2L1 |
| Q5BKY9 | 12 | Protein FAM | MAD2L1 |
| Q5VWQ0 | 4 | Lysine-specif | MAD2L1 |
| Q5W0B1 | 12.5 | RING finger | MAD2L1 |
| Q6NZI2 | 12 | Caveolae-ass | MAD2L1 |
| Q6PJG2 | 9 | ELM2 and SA | MAD2L1 |
| Q6ZNB6 | 11 | NF-X1-type z | MAD2L1 |
| Q70SY1 | 6 | Cyclic AMP-r | MAD2L1 |
| Q71UM5 | 93.5 | 40S ribosoma | MAD2L1 |
| Q7L7X3 | 9 | Serine/threo | MAD2L1 |
| Q7Z478 | 2 | ATP-depende | MAD2L1 |
| Q8NHQ9 | 7.5 | ATP-depende | MAD2L1 |
| Q8WWC4 | 10 | m-AAA prote | MAD2L1 |
| Q8WXI9 | 7.5 | Transcriptior | MAD2L1 |
| Q92667 | 8.5 | A-kinase anc | MAD2L1 |
| Q96CW1 | 10.5 | AP-2 comple | MAD2L1 |
| Q96EY4 | 38.5 | Translation r | MAD2L1 |
| Q96HE9 | 32 | Proline-rich p | MAD2L1 |
| Q96HS1 | 22 | Serine/threo | MAD2L1 |
| Q96KM6 | 6.5 | Zinc finger p | MAD2L1 |
| Q96L73 | 5 | Histone-lysin | MAD2L1 |
| Q96ME7 | 16.5 | Zinc finger p | MAD2L1 |
| Q96QE3 | 4 | ATPase fami | MAD2L1 |
| Q9BQ39 | 8 | ATP-depende | MAD2L1 |
| Q9BQ48 | 53 | 39S ribosoma | MAD2L1 |
| Q9BW60 | 10 | Elongation o | MAD2L1 |
| Q9BZF3 | 6 | Oxysterol-bir | MAD2L1 |
| Q9H0U4 | 86 | Ras-related | MAD2L1 |

| | | | |
|---|---|---|---|
| Q9H2Y7 | 4 | Zinc finger p | MAD2L1 |
| Q9H4L5 | 29 | Oxysterol-bir | MAD2L1 |
| Q9H8M2 | 10 | Bromodomai | MAD2L1 |
| Q9NPF2 | 12 | Carbohydrate | MAD2L1 |
| Q9NPG3 | 11 | Ubinuclein-1 | MAD2L1 |
| Q9NY93 | 17 | Probable ATI | MAD2L1 |
| Q9NZ01 | 19 | Very-long-ch | MAD2L1 |
| Q9P275 | 3 | Ubiquitin car | MAD2L1 |
| Q9P2N5 | 6.66666667 | RNA-binding | MAD2L1 |
| Q9UGP8 | 10 | Translocatio | MAD2L1 |
| Q9UHF7 | 2 | Zinc finger tr | MAD2L1 |
| Q9UIF8 | 1 | Bromodomai | MAD2L1 |
| Q9Y6J0 | 4 | Calcineurin-k | MAD2L1 |

| ID |
| --- |
| GO:0051988 |
| GO:0000776 |
| GO:0000777 |
| GO:0000778 |
| GO:0000942 |
| GO:0000940 |
| GO:0051987 |
| GO:0034501 |
| GO:0051382 |
| GO:0051315 |
| GO:0051455 |
| GO:0043515 |
| GO:0005828 |
| GO:1902425 |
| GO:0008608 |
| GO:0000941 |

| Ontology |
|---|
| regulation of attachment of spindle microtubules to kinetochore |
| kinetochore |
| condensed chromosome kinetochore |
| condensed nuclear chromosome kinetochore |
| condensed nuclear chromosome outer kinetochore |
| condensed chromosome outer kinetochore |
| positive regulation of attachment of spindle microtubules to kinetochore |
| protein localization to kinetochore |
| kinetochore assembly |
| attachment of mitotic spindle microtubules to kinetochore |
| attachment of spindle microtubules to kinetochore involved in homologous chromosome segregation |
| kinetochore binding |
| kinetochore microtubule |
| positive regulation of attachment of mitotic spindle microtubules to kinetochore |
| attachment of spindle microtubules to kinetochore |
| condensed nuclear chromosome inner kinetochore |

| ID | Ontology |
|---|---|
| GO:0007094 | mitotic spindle assembly checkpoint |
| GO:0090307 | mitotic spindle assembly |
| GO:0072686 | mitotic spindle |
| GO:0007052 | mitotic spindle organization |
| GO:1990023 | mitotic spindle midzone |
| GO:0060236 | regulation of mitotic spindle organization |
| GO:1990498 | mitotic spindle microtubule |
| GO:1901673 | regulation of mitotic spindle assembly |
| GO:0000070 | mitotic sister chromatid segregation |
| GO:0040001 | establishment of mitotic spindle localization |
| GO:0051256 | mitotic spindle midzone assembly |

| ID |
| --- |
| GO:0007094 |
| GO:0051988 |
| GO:0000776 |
| GO:0090307 |
| GO:0000777 |
| GO:0000778 |
| GO:0000942 |
| GO:0000940 |
| GO:0072686 |
| GO:0007052 |
| GO:0051987 |
| GO:0034501 |
| GO:0051382 |
| GO:0051315 |
| GO:0051455 |
| GO:0043515 |
| GO:0005828 |
| GO:1990023 |
| GO:0060236 |
| GO:1990498 |
| GO:1901673 |
| GO:1902425 |
| GO:0000070 |
| GO:0040001 |
| GO:0008608 |
| GO:0051256 |
| GO:0000941 |

| **Ontology** |
| --- |
| mitotic spindle assembly checkpoint |
| regulation of attachment of spindle microtubules to kinetochore |
| kinetochore |
| mitotic spindle assembly |
| condensed chromosome kinetochore |
| condensed nuclear chromosome kinetochore |
| condensed nuclear chromosome outer kinetochore |
| condensed chromosome outer kinetochore |
| mitotic spindle |
| mitotic spindle organization |
| positive regulation of attachment of spindle microtubules to kinetochore |
| protein localization to kinetochore |
| kinetochore assembly |
| attachment of mitotic spindle microtubules to kinetochore |
| attachment of spindle microtubules to kinetochore involved in homologous chromosome segregation |
| kinetochore binding |
| kinetochore microtubule |
| mitotic spindle midzone |
| regulation of mitotic spindle organization |
| mitotic spindle microtubule |
| regulation of mitotic spindle assembly |
| positive regulation of attachment of mitotic spindle microtubules to kinetochore |
| mitotic sister chromatid segregation |
| establishment of mitotic spindle localization |
| attachment of spindle microtubules to kinetochore |
| mitotic spindle midzone assembly |
| condensed nuclear chromosome inner kinetochore |

| ID | Ontology |
| --- | --- |
| GO:0005643 | nuclear pore |
| GO:0044615 | nuclear pore nuclear basket |
| GO:0017056 | structural constituent of nuclear pore |
| GO:0006999 | nuclear pore organization |
| GO:0051292 | nuclear pore complex assembly |
| GO:0031080 | nuclear pore outer ring |
| GO:0044613 | nuclear pore central transport channel |
